# Resurrection of plant disease resistance proteins via helper NLR bioengineering

**DOI:** 10.1101/2022.12.11.519957

**Authors:** Mauricio P. Contreras, Hsuan Pai, Muniyandi Selvaraj, AmirAli Toghani, David M. Lawson, Yasin Tumtas, Cian Duggan, Enoch Lok Him Yuen, Clare E. M. Stevenson, Adeline Harant, Abbas Maqbool, Chih-Hang Wu, Tolga O. Bozkurt, Sophien Kamoun, Lida Derevnina

## Abstract

Parasites counteract host immunity by suppressing helper NLR proteins that function as central nodes in immune receptor networks. Understanding the mechanisms of immunosuppression can lead to strategies for bioengineering disease resistance. Here, we show that a cyst nematode virulence effector binds and inhibits oligomerization of the helper NLR protein NRC2 by physically preventing intramolecular rearrangements required for activation. A single amino acid polymorphism at the binding interface between NRC2 and the inhibitor is sufficient for this helper NLR to evade immune suppression, thereby restoring the activity of multiple disease resistance genes. This points to a novel strategy for resurrecting disease resistance in crop genomes.

**One sentence summary:** A helper NLR is mutated to evade inhibition by a parasite effector.

## Introduction

The nucleotide binding and leucine-rich repeat (NLR) class of intracellular immune receptors are an important component of innate immunity in plants and animals. They mediate intracellular recognition of pathogens and subsequently initiate an array of immune responses in order to counteract infection (*1, 2*). NLRs can be activated by pathogen-secreted virulence proteins, termed effectors, which pathogens deliver into host cells to modulate host physiology (*2*). A hallmark of plant and animal NLR activation is their oligomerization into higher order immune complexes termed resistosomes or inflammasomes, respectively (*3-9*). These complexes initiate immune signaling via diverse mechanisms, often leading to a form of programmed cell death, termed hypersensitive response (HR) in plants or pyroptosis in animals (*10, 11*). Recent studies have reported NLR-like proteins mediating antiviral immunity and programmed cell death in prokaryotes via mechanisms analogous to those found in eukaryotic NLRs, suggesting that this is a conserved defense mechanism across all three domains of life (*12*). Remarkably, pathogen effectors can act both as triggers and suppressors of NLR-mediated immunity (*13*). In some cases, adapted pathogens deploy effectors which directly or indirectly interfere with NLR signaling to suppress immune activation (*12, 14-17*). However, the exact biochemical mechanisms by which pathogen effectors compromise NLR-mediated immunity to promote disease remain largely unknown. Moreover, whereas multiple strategies to bioengineer novel effector recognition specificities in NLRs have been proposed in recent years (*18*), approaches to mitigate the impact of effector-mediated immune suppression of NLRs are lacking.

NLRs belong to the signal ATPases with numerous domains (STAND) superfamily. They typically exhibit a tripartite domain architecture consisting of an N-terminal signaling domain, a central nucleotide binding domain and C-terminal superstructure forming repeats (*2*). The central domain, termed NB-ARC (nucleotide binding adaptor shared by APAF-1, plant R proteins and CED-4) in plant NLRs, is a hallmark of this protein family and plays a key role as a molecular switch, mediating conformational changes required for activation. NB-ARC domains consist of a nucleotide binding domain (NB), a helical domain (HD1), and a winged-helix domain (WHD) (*2, 19*). Diverse NLR activation and signaling strategies are found in nature. In some cases one NLR protein, termed a singleton, can mediate both elicitor perception and subsequent immune signaling (*20*). However, some NLRs can function as receptor pairs or in higher order configurations termed immune receptor networks (*13, 21*). In these cases, one NLR acts as a pathogen sensor, requiring a second helper NLR to initiate immune signaling. Such is the case in the solanaceous NRC immune receptor network, which is comprised of multiple sensor NLRs that require an array of downstream helper NLRs termed NRCs (NLRs required for cell death) to successfully initiate immune signaling. Upon perception of their cognate effectors, sensors in this network activate oligomerization of their downstream NRC helpers into a putative NRC resistosome, without stably forming part of the mature complex. This has been termed the activation-and release model (*4*). The NRC network can encompass up to half of the NLRome in some solanaceous plant species and plays a key role in mediating immunity against a variety of plant pathogens including oomycetes, bacteria, viruses, nematodes and insects (*15, 21*).

Plant and metazoan parasites have evolved effectors that interfere with host NLR signaling to promote disease. Parasite effectors can suppress NLR-mediated immunity indirectly by interfering with host proteins that act downstream of NLR signaling (*15, 17, 22, 23*), or directly by interacting with NLRs to inhibit their functions (*15, 16, 24*). One such example is the potato cyst nematode (*Globodera rostochiensis*) effector, SPRYSEC15 (SS15), which can suppress signaling mediated by *Nicotiana benthamiana* helper NLRs NRC2 and NRC3 and tomato (*Solanum lycopersicum*) helper NLR NRC1, by binding to their central NB-ARC domains (*15*) (**Fig. 1A, Fig. S1A**). In this study, we reasoned that mapping the binding interface between SS15 and its target helper NLRs would enable us to bioengineer NLR variants that evade pathogen suppression, therefore resurrecting the immune signaling activity of upstream sensors in the NRC network.

**Fig. 1:**
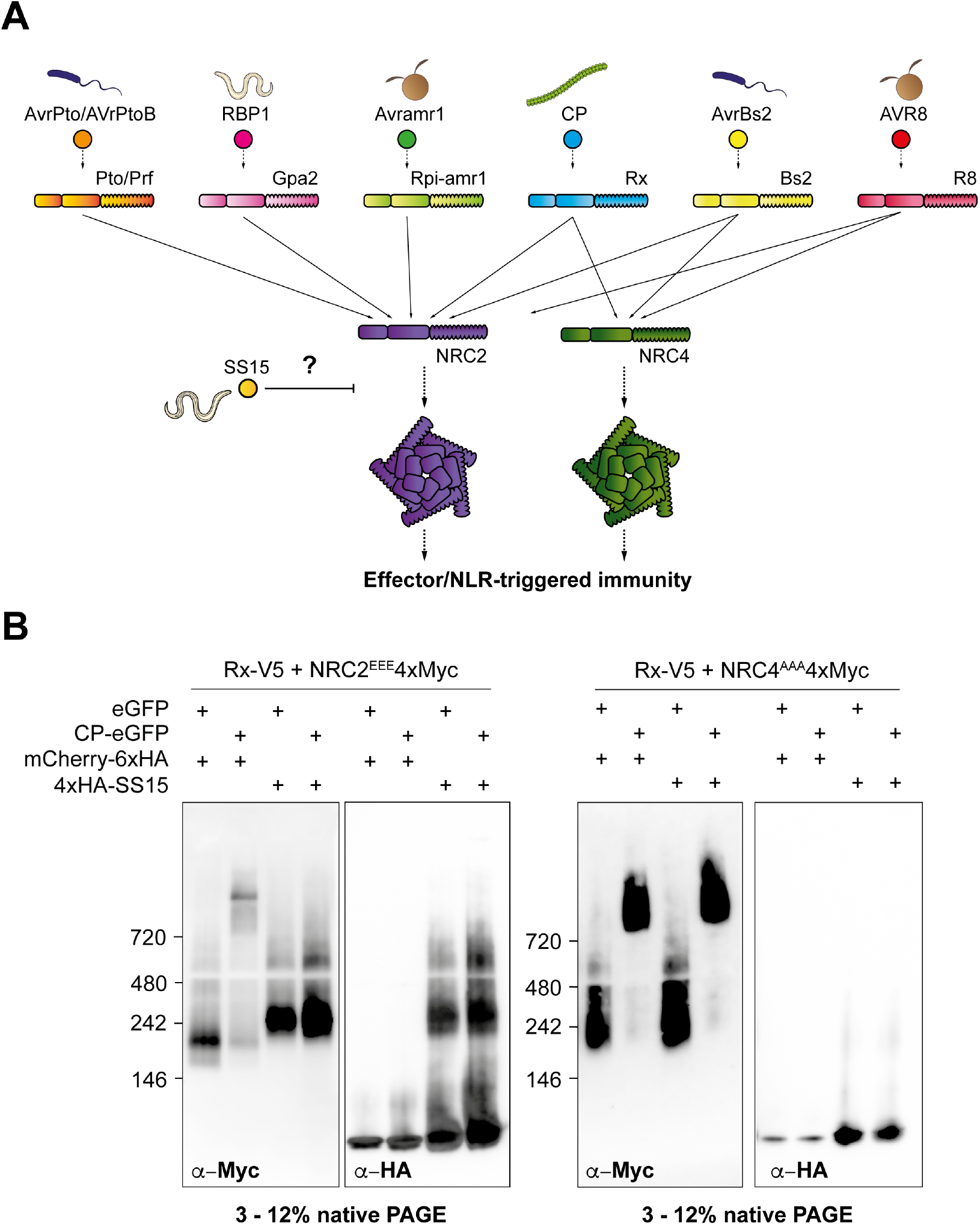
SS15 directly inhibits NRC2, but not NRC4 oligomerization. (**A**) Schematic representation of the NRC immune receptor network, consisting of multiple sensor NLRs (in blue) and their downstream helper NLRs, NRC2 and NRC4 (in purple and green, respectively). Effector-triggered activation of a sensor leads to downstream helper oligomerization and resistosome formation. The *G. rostochiensis* effector SS15 (in yellow) can directly bind to the NB-ARC domain of NRC2 but not NRC4, thereby inhibiting signaling by directly binding to the NB-ARC domain of this helper NLR. (**B**) BN-PAGE assays with inactive and activated Rx together with NRC2 or NRC4, in the absence or presence of SS15. C-terminally V5-tagged Rx and C-terminally 4xMyc-tagged NRC2^EEE^ or NRC4^AAA^ were co-expressed with either free GFP or C-terminally GFP-tagged PVX CP. These effector-sensor-helper combinations were co-infiltrated together with a 6xHA-mCherry fusion protein or with N-terminally 6xHA-tagged SS15. Total protein extracts were run in parallel on native and denaturing PAGE assays and immunoblotted with the appropriate antisera labelled below. Approximate molecular weights (kDa) of the proteins are shown on the left. Corresponding SDS-PAGE assays can be found in **Fig. S1**. The experiment was repeated three times with similar results.

## Results

First, we investigated how SS15 binding to NRC2 prevents immune signaling., notably whether SS15 prevents oligomerization of NRC following sensor NLR activation. To test this hypothesis, we transiently co-expressed NRC2 or NRC4 with their upstream sensor Rx and the effector SS15 in leaves of *nrc2/3/4* CRISPR KO *N. benthamiana* plants and leveraged previously established BN-PAGE-based readouts for NRC resistosome formation (*4*). For biochemical analyses, we used previously described NRC2 and NRC4 variants with mutations in their N-terminal MADA motifs (NRC2^EEE^ and NRC4^AAA^, respectively) which abolish cell death induction without compromising receptor activation, oligomerization, or localization (*4, 25, 26*). We activated the Rx-NRC system by co-expressing PVX CP-GFP or free GFP as an inactive control. In the absence of SS15, both NRC2 and NRC4 oligomerize upon effector-triggered activation mediated by their upstream sensor Rx. However, when SS15 is present, Rx/CP-activated NRC2 is unable to oligomerize and appears as a band of ∼240 kDa, which co-migrates with SS15. Inactive NRC2 co-expressed with SS15 also migrates as a band of ∼240 kDa, which is slower-migrating relative to inactive NRC2 in the absence of SS15, indicative of *in vivo* NRC2-SS15 complex formation (**Fig. 1B, Fig. S1**). We also observed that SS15 co-expression not only blocks NRC2 oligomerization but also prevents the previously reported shift of NRC2 from cytoplasm to PM as well as the formation of NRC2 PM-associated puncta upon Rx/CP activation (**Fig. S2**) (*4*). In contrast, NRC4 oligomerization is not affected in the presence of SS15, which is in line with previous findings that NRC4 immune signaling is not suppressed by SS15 (*15*). We conclude that SS15 can suppress immune signaling by acting as a direct proteinaceous inhibitor of NRC2, but not NRC4, by directly binding to its NB-ARC domain to block the formation of a signal-competent oligomeric resistosome.

Given that NRC4 retains the capacity to oligomerize in the presence of SS15, we leveraged this differential SS15 sensitivity between NRC2 and NRC4 to identify the domain that determines SS15 association and inhibition. We generated a series of NRC2-NRC4 chimeric proteins (**Fig. 2A-2B, Fig. S3**), which we subsequently assayed for SS15 association via *in planta* co-immunoprecipitation. We identified one chimeric variant of NRC4, carrying the HD1-1 region of NRC2 (NRC4^2HD1-1^), which gains association to SS15 (**Fig. 2C**). Unlike NRC4, NRC4^2HD1-1^ is susceptible to inhibition by SS15 and is unable to oligomerize and trigger cell death in the presence of SS15 (**Fig. 2D-2E, Fig. S3, Fig. S4**). We conclude that the HD1-1 region is important for association to SS15 and for the effector to directly inhibit NRC oligomerization and programmed cell death.

**Fig. 2:**
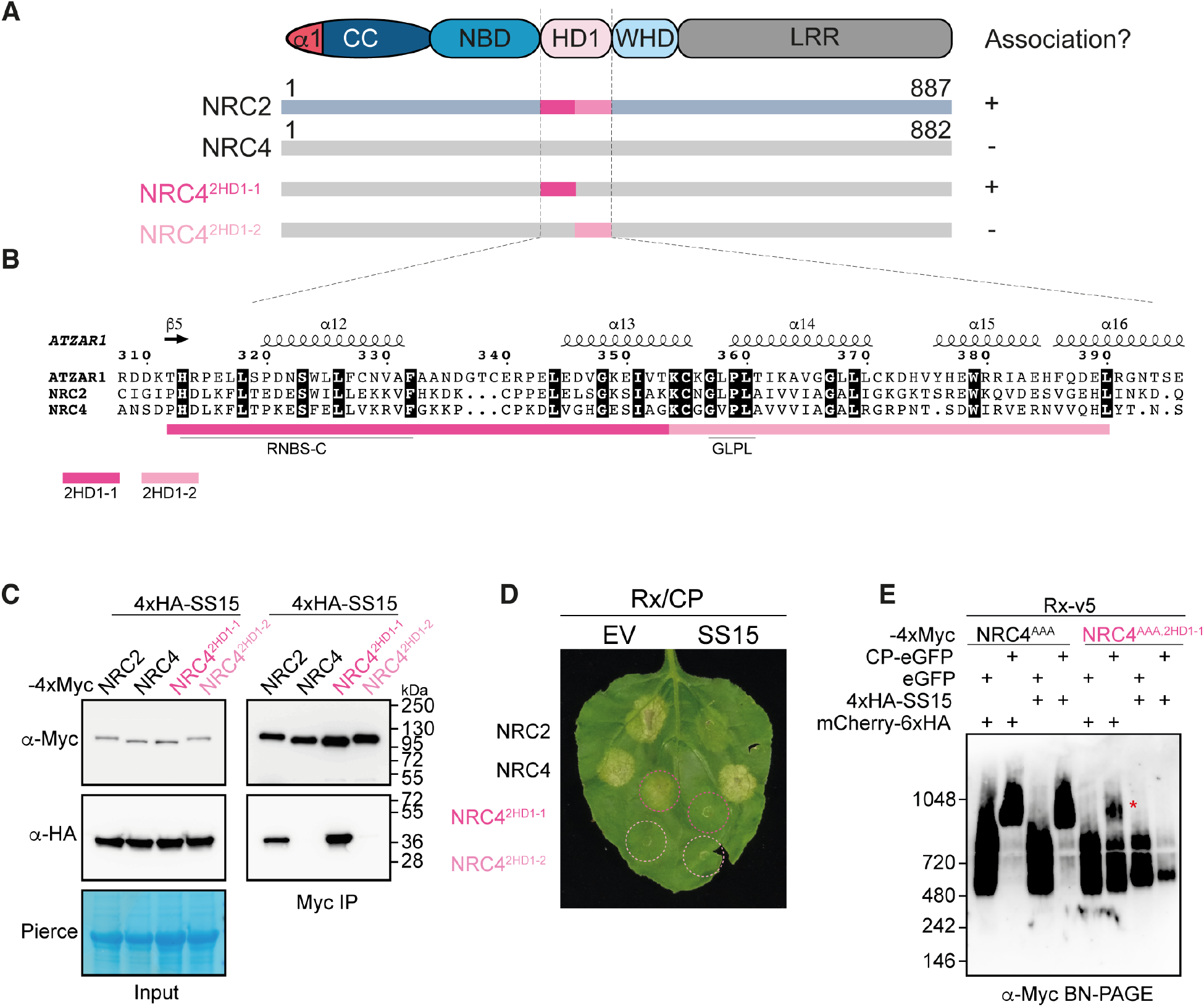
The HD1-1 region of NRC NB-ARC domains determines sensitivity to SS15. **(A)** Schematic representation of NRC domain architecture, highlighting regions within the NB-ARC domain swapped between NRC2-NRC4 chimeric proteins, namely NRC4^2HD1-1^ and NRC4^2HD1-2^. Association (+) or lack thereof (-) between these NLR immune receptors and SS15 determined by *in planta* co-immunoprecipitation is detailed on the right. (**B**) Close-up view of amino acid sequence alignment between AtZAR1, NRC2 and NRC4 focused on the HD1 region of NB-ARC domain. Predicted secondary structure is shown above the alignment. Well-characterized motifs within this region, such as RNBS-C and GLPL are underlined below the alignment. (**C**) Co-Immunoprecipitation (Co-IP) assays between SS15 and chimeric NRC2-NRC4 variants. C-terminally 4xMyc-tagged NRC proteins were transiently co-expressed with N-terminally 4xHA-tagged SS15. IPs were performed with agarose beads conjugated to Myc antibodies (Myc IP). Total protein extracts were immunoblotted with the appropriate antisera labelled on the left. Approximate molecular weights (kDa) of the proteins are shown on the right. Rubisco loading control was carried out using Ponceau stain (PS). The experiment was repeated three times with similar results. (**D**) Photo of representative leaves from *N. benthamiana nrc2/3/4* KO plants showing HR after co-expression of Rx and PVX CP with NRC2, NRC4 and the two NRC2-NRC4 chimeras, NRC4^2HD1-1^ and NRC4^2HD1-2^. These effector-sensor-helper combinations were co-expressed with a free mCherry-6xHA fusion protein (EV) or with N-terminally 4xHA-tagged SS15. The experiment consisted of 3 biological replicates with at least 6 technical replicates each. A quantitative analysis of the HR phenotypes can be found in **Fig. S4**. (**E**) BN-PAGE assay with inactive and activated Rx together with NRC4 or an NRC2-NRC4 chimeric protein in the absence or presence of SS15. C-terminally V5-tagged Rx and C-terminally 4xMyc-tagged NRC4^AAA^ or NRC4^AAA-2HD1-1^ were co-expressed with either free GFP or C-terminally GFP-tagged PVX CP. These effector-sensor-helper combinations were co-infiltrated together with a mCherry-6xHA fusion protein or with N-terminally 4xHA-tagged SS15. Total protein extracts were run in parallel on native and denaturing PAGE assays and immunoblotted with the appropriate antisera labelled on the left. Corresponding SDS-PAGE blots can be found in **Fig. S3**. Approximate molecular weights (kDa) of the proteins are shown on the left. The experiment was repeated three times with similar results.

To further define the interface between SS15 and NRC proteins, we crystallized SS15 in complex with the NB-ARC domain of SlNRC1, a tomato NRC that is inhibited by the nematode effector. We subsequently solved the structure using X-ray diffraction data collected to 4.5 Å resolution (**Fig. 3A, Fig. S5, Table S3**), which allowed us to determine that SS15 binds to a loop in the HD1-1 region of NRCs which connects the NB domain to the HD1 and WHD domains. This provides orthogonal evidence that the SS15-NRC interactions are mediated by this region as shown with the chimera experiments (**Fig. 2**). This loop was previously shown to act as a “hinge”, allowing the NB domain to rotate relative to the HD1 and WHD domains following activation (**Fig. S5, Movie S1**) (*27*). We propose that SS15 prevents conformational changes that are critical for NLR activation by binding and immobilizing the NB-HD1 hinge.

**Fig. 3:**
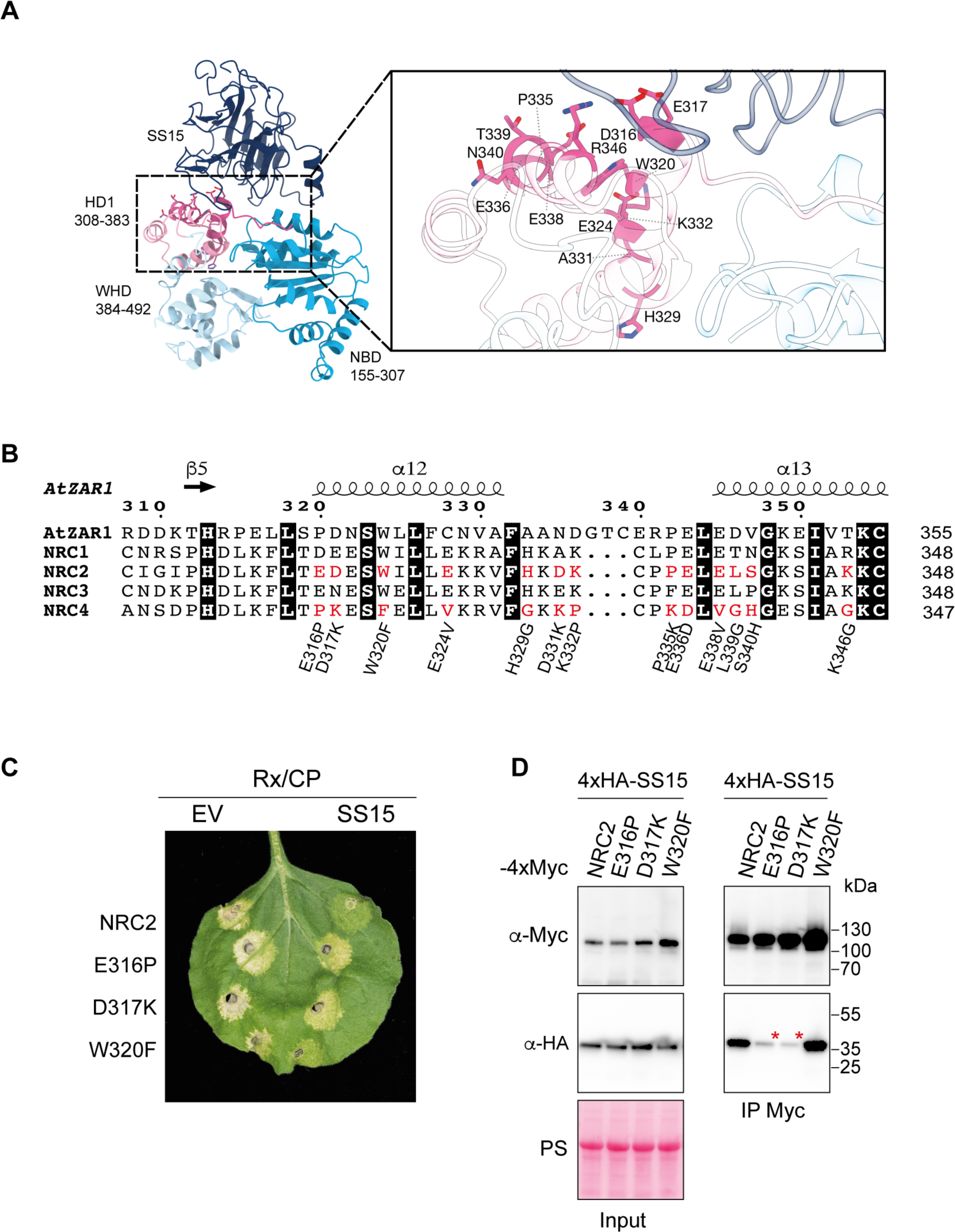
SS15-NRC binding interface enables bioengineering NRC2 variants that evade suppression. **(A)** Crystal structure of the SS15-SlNRC1^NB-ARC^ complex. The NB, HD1 and WHD domains of SlNRC1^NB-ARC^ are shown in cyan, pale blue and magenta, respectively; SS15 is in dark blue. The inset displays a close-up of the interface between SS15 and the HD1 domain of SlNRC1^NB-ARC^ with the residues corresponding to those selected for mutagenesis in NRC2 highlighted in stick representation. See **Table S3** for a summary of data collection and processing statistics. (**B**) Alignment of HD1-1 region of AtZAR1, SlNRC1, NRC2, NRC3, and NRC4. Candidate residues (highlighted in red) were shortlisted based on the interface identified in the co-crystal structure of SS15 and the SlNRC1 NB-ARC domain, as well as being conserved in SlNRC1, NRC2 and NRC3 but not NRC4 and AtZAR1. 13 NRC2 variants were generated by mutating individual candidate positions to the corresponding amino acid in NRC4 (detailed underneath the alignment). (**C**) Photo of representative leaves from *N. benthamiana nrc2/3/4* KO plants showing HR after co-expression of Rx and PVX CP with NRC2, or different NRC2 variants generated. These effector-sensor-helper combinations were co-expressed with a free mCherry-6xHA fusion protein (EV) or with N-terminally 4xHA-tagged SS15. The experiment consisted of 3 biological replicates. A quantitative analysis of HR phenotypes can be found in **Fig. S7**. (**D**) Co-Immunoprecipitation (Co-IP) assays between SS15 and NRC2 variants. C-terminally 4xMyc-tagged NRC2 variants were transiently co-expressed with N-terminally 4xHA-tagged SS15. IPs were performed with agarose beads conjugated to Myc antibodies (Myc IP). While only data for E316P, D317K and W320F NRC2 variants is shown, additional variants were also tested and can be found in **Fig S6**. Total protein extracts were immunoblotted with appropriate antisera labelled on the left. Approximate molecular weights (kDa) of the proteins are shown on the right. Rubisco loading control was carried out using Ponceau stain (PS). The experiment was repeated three times with similar results.

Given that SS15 suppresses cell death induction mediated by SlNRC1, NRC2 and NRC3 but not NRC4 or other well characterized NLRs such as ZAR1 (**Fig. S1**) (*15*), we leveraged the high degree of conservation that is characteristic of plant NB-ARC domains to narrow down residues within the binding interface that underpin this interaction. We shortlisted residues within the HD1-1 region that are similar in SlNRC1, NRC2 and NRC3 but different in NRC4 or AtZAR1 (**Fig. 3B**). Combining information from the co-crystal structure and the alignments allowed us to select 13 candidate residues to test by mutagenesis in NRC2 (**Fig. 3B**). We mutated each of these residues to the corresponding amino acid found in NRC4 and screened these NRC2 variants for susceptibility to SS15 inhibition in cell death assays. This revealed two variants, NRC2^E316P^ and NRC2^D317K^ which triggered cell death when activated by Rx/CP and were no longer inhibited by SS15 (**Fig. 3C, Fig. S6, Fig. S7**). We also tested all 13 single amino acid mutants for association with SS15 by *in planta* co-immunoprecipitation and found that NRC2^E316P^ and NRC2^D317K^ exhibited reduced association with SS15 relative to NRC2 (**Fig. 3D, Fig. S6**), which is in line with the observation that SS15 is not able to suppress these variants (**Fig. 3C**). We conclude that the E316 and D317 residues are critical for SS15-mediated inhibition of NRC2 and that mutating these residues to their equivalent amino acid in NRC4 allows Rx/CP-activated NRC2 to evade SS15 association and inhibition.

We next tested whether these two SS15-evading variants of NRC2 could restore functionality of NRC2-dependent sensor NLRs that are suppressed by the parasite effector. We tested this by performing complementation assays with NRC2^E316P^ and NRC2^D317K^ in *nrc2/3/4* CRISPR KO *N. benthamiana* plants. We activated the NRC2 variants with a panel of agronomically important sensor NLRs mediating resistance to diverse pathogens, including the potato cyst nematode R gene Gpa2 (an allele of Rx), as well as other well characterized oomycete and bacterial resistance proteins. Remarkably, NRC2^D317K^ evaded SS15 inhibition with all tested sensor NLRs restoring their capacity to activate immune signaling (**Fig 4A, Fig. S8, Fig. S9**). In contrast, NRC2^E316P^ could evade SS15 suppression when activated by Rx, but not when activated by other sensors. We therefore selected NRC2^D317K^ for follow-up biochemical studies, using BN-PAGE-based assays. Unlike NRC2, activated NRC2^D317K^ oligomerized even in the presence of SS15 and did not form an *in vivo* complex with the inhibitor (**Fig. 4B**). We conclude that NRC2^D317K^ can fully evade SS15-mediated immune suppression, retaining the capacity to oligomerize and mediate cell death when activated by multiple agronomically important sensor NLRs.

**Fig. 4:**
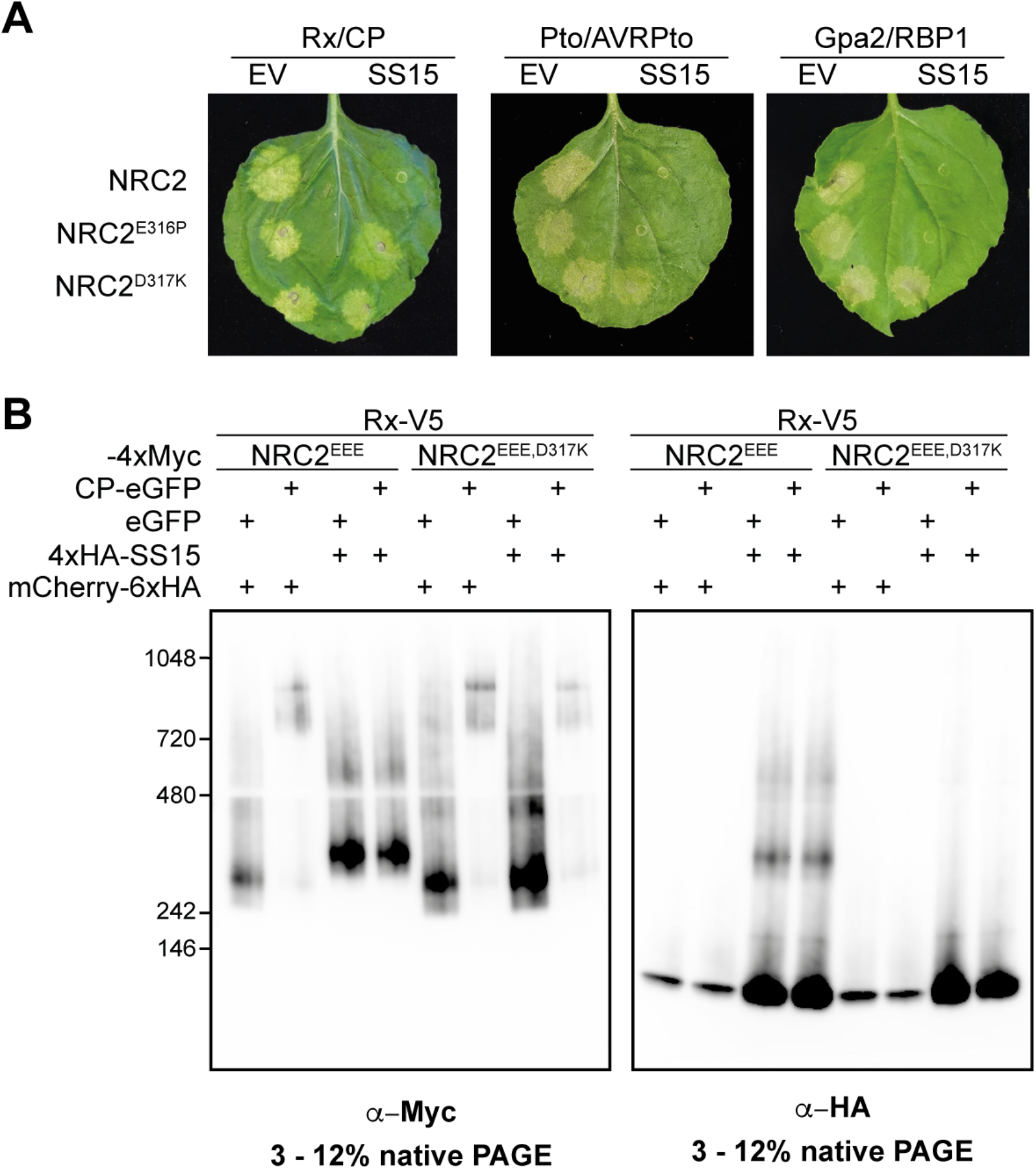
NRC2^D317K^ helper restores immune signaling of multiple disease resistance genes in the presence of the effector SS15. **(A)** Photo of representative leaves from *N. benthamiana nrc2/3/4* KO plants showing HR after co-expression of Rx/CP, Pto/AVRPto or Gpa2/RBP1 together with NRC2, NRC2^E316P^ or NRC2^D317K^ in the absence or presence of SS15. The experiment consisted of 3 biological replicates. A quantitative analysis of HR phenotypes can be found in **Fig. S8**. (**B**) BN-PAGE assays with inactive and activated Rx together with NRC2 or NRC2^D317K^, in the absence or presence of SS15. C-terminally V5-tagged Rx and C-terminally 4xMyc-tagged NRC2^EEE^ or NRC2^EEE-D317K^ were co-expressed with either free GFP or C-terminally GFP-tagged PVX CP. These effector-sensor-helper combinations were co-infiltrated together with a 6xHA-mCherry fusion protein or with N-terminally 4xHA-tagged SS15. Total protein extracts were run in parallel on native and denaturing PAGE assays and immunoblotted with the appropriate antisera labelled below. Approximate molecular weights (kDa) of the proteins are shown on the left. Corresponding SDS-PAGE blots can be found in **Fig. S9**. The experiment was repeated three times with similar results.

## Conclusion

Our study reveals how a parasite effector has evolved as an inhibitor of a helper NLR by directly binding its NB-ARC domain to prevent resistosome formation and immune signaling. By binding and immobilizing a critical hinge loop in the HD1 region of the NB-ARC, SS15 restricts movement of the NB domain relative to the HD1 and WHD domains, preventing immune receptor activation. Remarkably, while SS15 can bind and inhibit NRC2, it cannot bind or inhibit the NRC2 paralog NRC4. We exploited NRC4’s resilience to SS15 inhibition by making chimeric NRC2-NRC4 variants which, together with structural information, helped us identify the inhibitor binding interface. Mutational studies of this interface allowed us to generate a single amino acid variant of NRC2 (NRC2^D317K^) that evades SS15 inhibition without compromising receptor signaling capacity. This NRC2^D317K^ variant can now support signaling by any NRC2-dependent sensor even in the presence of SS15.

The existence of plant parasite secreted NLR inhibitors suggests that suppressed resistance genes may occur in crop genomes. Leveraging the approach detailed in this study, it may be possible to resurrect cryptic or defeated resistance proteins to enhance disease resistance. Moreover, considering that multiple sensors can signal through the same downstream helper, applying this approach to helper NLRs holds potential to simultaneously resurrect multiple upstream sensor NLRs. Importantly, the single amino acid NRC2 variants we identified could in theory be generated in-locus using gene editing technologies in agronomically important crop species, making deployment of this technology viable in countries where transgenic approaches are not feasible. Our work describes a novel approach to achieve robust immunity by engineering NLRs that avoid parasite suppression. This could in theory be applied to other plant, metazoan or even prokaryotic NLR immune receptors that are directly targeted by parasite effectors (*12, 15, 16, 24*). Combined with recent advances in NLR engineering to achieve novel pathogen recognition specificities (*18, 28*), this technology holds the potential to unlock a new era in disease resistance breeding.

## Materials and Methods

### Plant growth conditions

Wild-type and *nrc2/3/4* CRISPR mutant *Nicotiana benthamiana* lines were grown in a controlled environment growth chamber with a temperature range of 22 to 25 ºC, humidity of 45% to 65% and a 16/8-hour light/dark cycle.

### Plasmid construction

We used the Golden Gate Modular Cloning (MoClo) kit (*29*) and the MoClo plants part kit (*30*) for cloning. All vectors used were generated with these kits unless otherwise stated. Cloning design and sequence analysis were done using Geneious Prime (v2021.2.2; https://www.geneious.com). Plasmid construction is described in more detail in **Table S1**.

### Cell death assays by agroinfiltration

Proteins of interest were transiently expressed in *N. benthamiana* according to previously described methods (*31*). Briefly, leaves from 4–5-week-old plants were infiltrated with suspensions of *Agrobacterium tumefaciens* GV3101 pM90 strains transformed with expression vectors coding for different proteins indicated. Final OD_600_ of all *A. tumefaciens* suspensions were adjusted in infiltration buffer (10 mM MES, 10 mM MgCl_2,_ and 150 μM acetosyringone (pH 5.6)). Final OD_600_ used for each construct is described in **Table S2**.

### Extraction of total proteins for BN-PAGE and SDS-PAGE assays

Four to five-week-old *N. benthamiana* plants were agroinfiltrated as described above with constructs of interest and leaf tissue was collected 3 days post agroinfiltration for NRC2 and 2 days post agroinfiltration for NRC4. Final OD_600_ used for each construct is described in **Table S2**. BN-PAGE was performed using the Bis-Tris Native PAGE system (Invitrogen) according to the manufacturer’s instructions, as described previously (*31*). Leaf tissue was ground using a Geno/Grinder tissue homogenizer. For NRC2, GTMN extraction buffer was used (10% glycerol, 50 mM Tris-HCl (pH 7.5), 5 mM MgCl_2_ and 50 mM NaCl) supplemented with 10 mM DTT, 1x protease inhibitor cocktail (SIGMA) and 0.2% Nonidet P-40 Substitute (SIGMA). For NRC4, GHMN buffer (10% glycerol, 50 mM HEPES (pH 7.4), 5 mM MgCl_2_ and 50 mM NaCl) buffer supplemented with 10mM DTT, 1x protease inhibitor cocktail (SIGMA) and 1% Digitonin (SIGMA) was used for extraction. Samples were incubated in extraction buffer on ice for 10 minutes with short vortex mixing every 2 minutes. Following incubation, samples were centrifuged at 5,000 x*g* for 15 minutes and the supernatant was used for BN-PAGE and SDS-PAGE assays.

### BN-PAGE assays

For BN-PAGE, samples extracted as detailed above were diluted as per the manufacturer’s instructions by adding NativePAGE 5% G-250 sample additive, 4x Sample Buffer and water. After dilution, samples were loaded and run on Native PAGE 3%-12% Bis-Tris gels alongside either NativeMark unstained protein standard (Invitrogen) or SERVA Native Marker (SERVA). The proteins were then transferred to polyvinylidene difluoride membranes using NuPAGE Transfer Buffer using a Trans-Blot Turbo Transfer System (Bio-Rad) as per the manufacturer’s instructions. Proteins were fixed to the membranes by incubating with 8% acetic acid for 15 minutes, washed with water and left to dry. Membranes were subsequently re-activated with methanol to correctly visualize the unstained native protein marker. Membranes were immunoblotted as described below.

### SDS-PAGE assays

For SDS-PAGE, samples were diluted in SDS loading dye and denatured at 72 ºC for 10 minutes. Denatured samples were spun down at 5,000 x*g* for 3 minutes and supernatant was run on 4%-20% Bio-Rad 4%-20% Mini-PROTEAN TGX gels alongside a PageRuler Plus prestained protein ladder (Thermo Scientific). The proteins were then transferred to polyvinylidene difluoride membranes using Trans-Blot Turbo Transfer Buffer using a Trans-Blot Turbo Transfer System (Bio-Rad) as per the manufacturer’s instructions. Membranes were immunoblotted as described below.

### Immunoblotting and detection of BN-PAGE and SDS-PAGE assays

Blotted membranes were blocked with 5% milk in Tris-buffered saline plus 0.01% Tween 20 (TBS-T) for an hour at room temperature and subsequently incubated with desired antibodies at 4 ºC overnight. Antibodies used were anti-GFP (B-2) HRP (Santa Cruz Biotechnology), anti-HA (3F10) HRP (Roche), anti-Myc (9E10) HRP (Roche), and anti-V5 (V2260) HRP (Roche), all used in a 1:5000 dilution in 5% milk in TBS-T. To visualize proteins, we used Pierce ECL Western (32106, Thermo Fisher Scientific), supplementing with up to 50% SuperSignal West Femto Maximum Sensitivity Substrate (34095, Thermo Fishes Scientific) when necessary. Membrane imaging was carried out with an ImageQuant LAS 4000 or an ImageQuant 800 luminescent imager (GE Healthcare Life Sciences, Piscataway, NJ). Rubisco loading control was stained using Ponceau S (Sigma).

### Co-Immunoprecipitation assays

CoIP assays were performed as described previously (*32*). Total soluble protein was extracted as described above from leaves of *N. benthamiana* 3 days post agro-infiltration using GTEN buffer (10% Glycerol, 25 mM Tris-HCl (pH 7.5), 1mM EDTA, 150 mM NaCl) supplemented with 2% (w/v) polyvinylpolypyrrolidone, 10 mM DTT and 1x protease inhibitor cocktail (SIGMA), 0.3% IGEPAL (SIGMA). OD_600_ used can be found in **Table S2**. Protein extracts were filtered using Minisart 0.45 μm filter (Sartorius Stedim Biotech, Goettingen, Germany). Part of the extract was set aside prior to immunoprecipitation and used for SDS-PAGE as described above. These were the inputs. 1.4 ml of the remaining filtered total protein extract was mixed with 30 μl of anti-c-Myc agarose beads (A7470, SIGMA) and incubated end over end for 90 minutes at 4 ºC. Beads were washed 5 times with immunoprecipitation wash buffer (GTEN extraction buffer with 0.3% v/v IGEPAL (SIGMA)) and resuspended in 60 μl of SDS loading dye. Proteins were eluted from beads by heating for 10 minutes at 72 ºC. Immunoprecipitated samples were used for SDS-PAGE and immunoblotted as described above and compared to the inputs.

### Confocal microscopy

Three to four-week-old plants were agroinfiltrated as described above with constructs of interest. Final OD_600_ used for each construct is described in **Table S2**. Leaf tissue was prepared for imaging by sectioning of desired area surrounding an infiltration spot using a cork borer size 4, and were mounted, live, in wells containing dH2O made in Carolina Observation Gel to enable diffusion of gasses. The abaxial of the leaf tissue was imaged using a Leica SP8 with 40x water immersion objective. Laser excitations for fluorescent proteins were used as described previously (*33*), namely 488 nm (Argon) for GFP, 561/594 nm (Diode) for RFP and 405 nm (Diode) for BFP.

### Membrane enrichment assays

Membrane enrichment was carried out by slightly modifying a previously described protocol (*34*). In brief, leaf material was ground to fine powder using liquid nitrogen and 2x volume of extraction buffer was added. Extraction buffer consisted of 0.81M sucrose, 5% (v/v) glycerol, 10 mM EDTA, 10 mM EGTA, 5mM KCl, and 100 mM Tris-HCl (pH 7.5) supplemented with 5 mM DTT, 1% Sigma Plant Protease Inhibitor Cocktail, 1 mM PMSF and 0.5% PVPP. After addition of the buffer, the samples were vortexed for a minute and the cell debris was cleared out by two subsequent centrifugation steps at 1000 xg for 5 min. The supernatant was diluted 1:1 using distilled water and an aliquot of the supernatant was separated as the total fraction (T). The remaining supernatant (200-300 μL) was further centrifuged at 21,000 xg for 90 min at 4°C. This centrifugation yielded the supernatant (soluble fraction, S) and membrane enriched pellet (membrane fraction, M). After separating the soluble fraction, the pellet was resuspended in diluted extraction buffer (without PVPP). All the fractions were diluted with SDS loading dye, and proteins were denatured by incubating at 50°C for 15 min. Western blotting was performed as previously described following SDS-PAGE. Endogenous plasma membrane ATPase was detected using anti-H+ATPase (AS07 260) antibody (Agrisera) as a marker to show the success of membrane enrichment.

### Heterologous protein production and purification from *E. coli*

Heterologous production and purification of SS15 was performed as previously described (*32*). Recombinant SS15 protein (lacking signal peptide) was expressed by cloning in pOPIN-S3C plasmid, with an N-terminal tandem 6xHis-SUMO followed by a 3C protease cleavage site. pOPIN-S3C:SS15 was transformed into *E. coli* SHuffle cells. 8L of these cells were grown at 30 ºC in autoinduction media (*35*) to an OD600 of 0.6 to 0.8 followed by overnight incubation at 18 ºC and harvested by centrifugation. Pelleted cells were resuspended in 50 mM Tris HCl (pH 8), 500 mM NaCl, 50 mM Glycine, 5% (vol/vol) glycerol and 20 mM imidazole (buffer A) supplemented with cOmplete EDTA-free protease inhibitor tablets (Roche) and lysed by sonication. The clarified cell lysate was applied to a Ni^2+^-NTA column connected to an AKTA pure system. 6xHis-SUMO-SS15 was step eluted with elution buffer (buffer A with 500 mM imidazole) and directly injected onto a Superdex 200 26/600 gel filtration column pre-equilibrated with buffer B (20 mM HEPES (pH 7.5), 150 mM NaCl). The fractions containing 6xHis-SUMO-SS15 were pooled, and the N-terminal 6xHis-SUMO tag was cleaved by addition of 3C protease (10 µg/mg of fusion protein), incubating overnight at 4ºC. Cleaved SS15 was further purified using a Ni^2+^-NTA column, this time collecting the flow through to separate the cleaved tag from the SS15 protein. Un-tagged SS15 was further purified by another round of gel filtration as described above. The concentration of protein was judged by absorbance at 280 nm (using a calculated molar extinction coefficient of 35920 M^-1cm-1^ for SS15).

Heterologous production and purification of SlNRC1^NB-ARC^ was performed as previously described (*36*). Recombinant SlNRC1^NB-ARC^ protein was also expressed cloning in pOPIN-S3C plasmid as described above. pOPIN-S3C:SlNRC1^NB-ARC^ was transformed into *E. coli* Lemo21 (DE3) cells. 8L of these cells were grown at 37 ºC in autoinduction media (*35*) to an OD600 of 0.6 to 0.8 followed by overnight incubation at 18 ºC and harvested by centrifugation. Pelleted cells were resuspended in 50 mM Tris HCl (pH 8), 500 mM NaCl, 50 mM Glycine, 5% (vol/vol) glycerol and 20 mM imidazole (buffer A) supplemented with cOmplete EDTA-free protease inhibitor tablets (Roche) and lysed by sonication. The clarified cell lysate was applied to a Ni^2+^-NTA column connected to an AKTA pure system. 6xHis-SUMO-SlNRC1^NB-ARC^ was step eluted with elution buffer (buffer A with 500 mM imidazole) and directly injected onto a Superdex 200 26/600 gel filtration column pre-equilibrated with buffer B (20 mM HEPES (pH 7.5), 150 mM NaCl). The fractions containing 6xHis-SUMO-SlNRC1^NB-ARC^ were pooled, and the N-terminal 6xHis-SUMO tag was cleaved by addition of 3C protease (10 µg/mg of fusion protein), incubating overnight at 4ºC. Cleaved SlNRC1^NB-ARC^ was further purified using a Ni^2+^-NTA column, this time collecting the flow through to separate the cleaved tag from the SlNRC1^NB-ARC^ protein. Un-tagged SlNRC1^NB-ARC^ was further purified by another round of gel filtration as described above. The concentration of protein was judged by absorbance at 280 nm (using a calculated molar extinction coefficient of 63370 M^-1cm-1^ for SlNRC1^NB-ARC^).

To obtain SlNRC1^NB-ARC^ in complex with SS15, both proteins were incubated in a 1:1 molar ratio overnight at 4 ºC and subjected to gel filtration on a Superdex 200 26/600 gel filtration column as described above. The fractions containing SlNRC1^NB-ARC^ in complex with SS15 were pooled, concentrated to 10-15 mg/ml and subsequently used for crystallisation screens.

### Crystallisation, data collection, and structure solution

Crystallisation screens were performed at 18°C using the sitting-drop vapour diffusion technique. Drops composed of 0.3 μL of protein solution and 0.3 μL of reservoir solution were set up in MRC 96-well crystallisation plates (Molecular Dimensions), which were dispensed using an Oryx Nano or an Oryx8 robot (Douglas Instruments). Crystal growth was monitored using a Minstrel Desktop Crystal Imaging System (Rikagu). Suitable crystals grew after 72 hrs in a Morpheus screen crystallisation condition containing 0.1M MES buffer (pH 6.5), 10% (w/v) PEG8000 and 20% (v/v) ethylene glycol (Molecular Dimensions) and were harvested by flash-cooling in liquid nitrogen using LithoLoops (Molecular Dimensions). X-ray diffraction data were collected at the Diamond Light Source (Didcot, UK) on beamline I03 using an Eiger2 XE 16M pixel array detector (Dectris) with crystals maintained at 100 K by a Cryojet cryocooler (Oxford Instruments).

X-ray data were integrated, and scaled using XDS (*37*), as implemented through the XIA2 (*38*) pipeline, and then merged using AIMLESS (*39*), via the CCP4i2 graphical user interface (*40*). The NRC1^NB-ARC^-SS15 complex crystallised in space group *P*6_1_ with cell parameters *a*=*b*=128.6, *c*=170.7 Å, and the best crystal yielded diffraction data to 4.5 Å resolution (see **Table S3** for a summary of data collection and refinement statistics). Given the small size of the dataset, we assigned 10% of the data (883 unique reflections) for the *R*_free_ calculation, to give a more statistically meaningful metric. The crystal structure of NRC1^NB-ARC^ alone was already available (PDB 6S2P), but there was no experimentally determined structure for SS15. Thus, we made use of AlphaFold2-multimer (*41*), as implemented through Colabfold (https://colab.research.google.com/github/sokrypton/ColabFold/blob/main/AlphaFold2.ipynb) (*42*) to generate structural predictions for the complex. There was very good sequence coverage for both proteins and the five independent models of the individual components were closely similar. The pLDDT scores were generally good (e.g. averages of 82 and 75 for NRC1^NB-ARC^ and SS15 models, respectively, from the rank 1 predictions). However, the relative placement of the two components varied across the five models and the corresponding PAE scores indicated very low confidence in these predictions. A comparison of the five NRC1^NB-ARC^ models with the known crystal structure showed a good agreement (e.g. superposition of the rank 1 model gave an rmsd of 1.77 Å). Given that the AlphaFold2 (AF2) model provided starting coordinates for several loops missing from the crystal structure, we decided to use this model in molecular replacement. Templates for both components were prepared using the “Process Predicted Models” CCP4i2 task, which removed low confidence regions (based on pLDDT) and converted the pLDDT scores in the *B*-factor field of the PDB coordinate files to pseudo *B*-factors. Analysis of the likely composition of the asymmetric unit (ASU) suggested that it contained two copies of each component, giving an estimated solvent content of ∼67%. PHASER (*43*) was able to place the four chains within the ASU, although the second SS15 domain required manual repositioning with respect to its neighbouring NRC1 domain in order to avoid a number of clashes and improve the fit to the density. This was achieved using COOT (*44*) and guided by the arrangement of the other NRC1-SS15 complex (**Fig. S4**). The structure was then subjected to jelly body refinement in REFMAC5 (*45*) using ProSMART restraints (*46*) generated from the AF2 models, giving *R*_work_ and *R*_free_ values of 0.357 and 0.401, respectively, to 4.5 Å resolution.

Now it was possible to generate more complete models for the components by superposing the original unprocessed AF2 models and trimming these with reference to the improved electron density. Furthermore, a significant region of positive difference density was present at the cores of both NRC1 domains, which corresponded to the ADP in the crystal structure; thus we incorporated ADP into the model. Due to the low resolution of the dataset, only very limited rebuilding was possible in COOT, where Geman-McClure and Ramachandran restraints were used to maintain good stereochemical parameters. After several cycles of restrained refinement in REFMAC5 and editing in COOT, a reasonable model was obtained with *R*_work_ and *R*_free_ values of 0.258 and 0.298, respectively. However, there remained a region of positive difference density near the N-termini of both SS15 domains that we could not adequately explain. At this point we re-ran the AlphaFold2-multimer predictions, but this time with one copy of the complex taken from the crystal structure as a reference template. Although the new predictions did not produce complexes that were consistent with the X-ray data, and the models for the individual components did not appear to be noticeably improved based on pLDDT scores, we used them as starting points to rebuild the X-ray structure.

Significantly, for several models, the N-terminal region of SS15 adopted conformations that partially accounted for the region of positive difference density electron density, and this could be improved by careful rebuilding and refinement. This “AlphaFold recycling” procedure was repeated a further two times before finalising the structure, which included residues 153-494 for SlNRC1 (numbered with respect to full length protein) and residues 18-223 for SS15, where residues 33-43 in both copies of the latter formed α-helices that occupied the regions of positive difference density observed earlier. For the last refinement in REFMAC5, the following options were used: ProSMART restraints generated from the latest AF2 models, overall *B*-factor refinement with TLS restraints (where each protein chain was treated as a separate domain), and non-crystallographic symmetry restraints. The final model gave *R*_work_ and *R*_free_ values of 0.237 and 0.275, respectively, to 4.5 Å resolution (see **Table S3** for a summary of refinement statistics). All structural figures were prepared using ChimeraX (*47*) and PyMOL (*48*).

## Supplemental Data

**Fig. S1:**
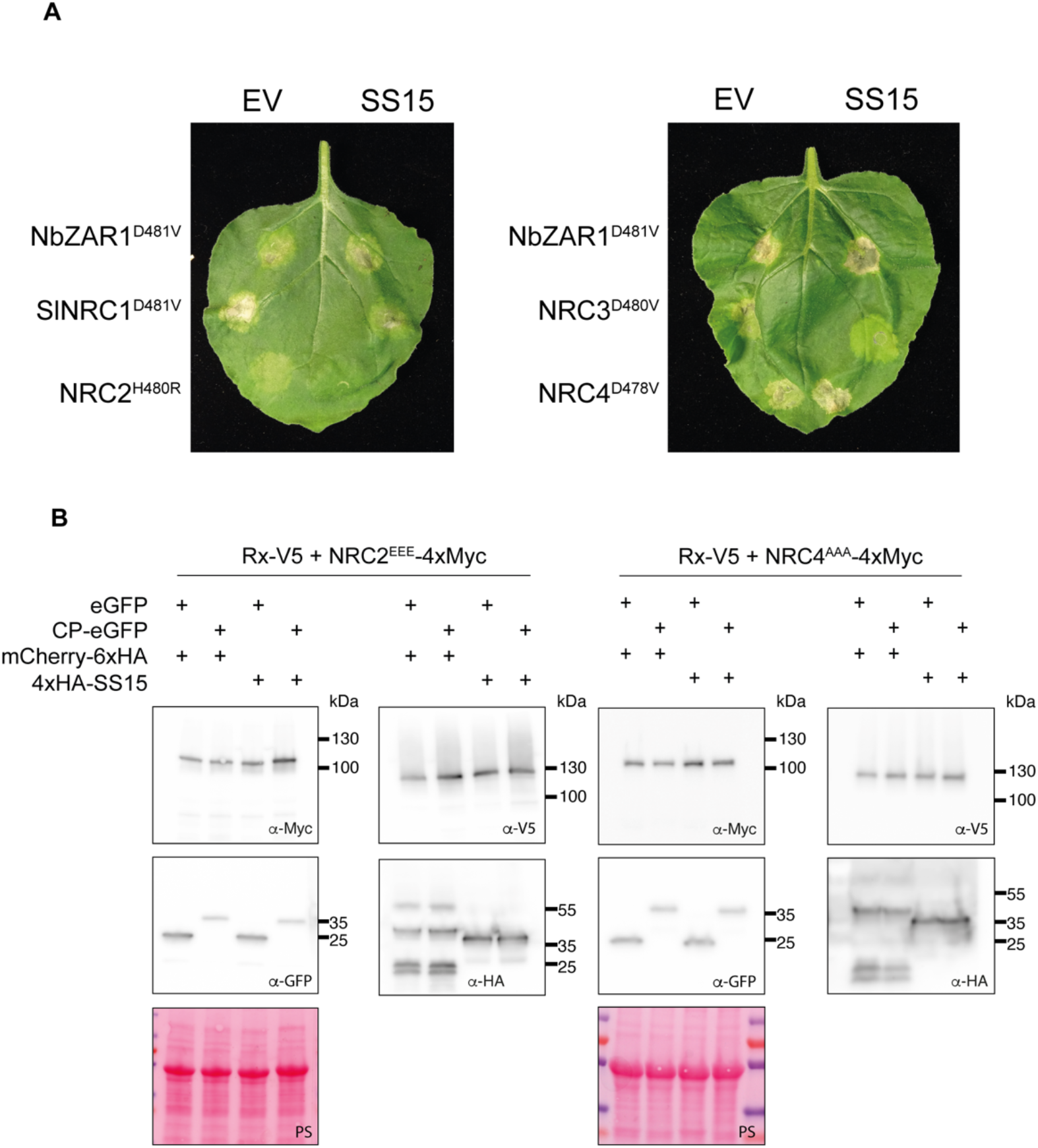
SS15 suppresses cell death mediated by SlNRC1, NRC2 and NRC3 but not NRC4 or NbZAR1. (**A**) Photo of representative leaves from *N. benthamiana nrc2/3/4* KO plants showing HR after co-expression of various autoactive NLR variants with a free mCherry-6xHA fusion protein (EV) or with N-terminally 4xHA-tagged SS15. Images are representative of (**B**) SDS-PAGE accompanying BN-PAGE shown in **Fig. 1B**. Total protein extracts were immunoblotted with the appropriate antisera labelled on the left. Approximate molecular weights (kDa) of the proteins are shown on the right. Rubisco loading control was carried out using Ponceau stain (PS). The experiment was repeated three times with similar results.

**Fig. S2:**
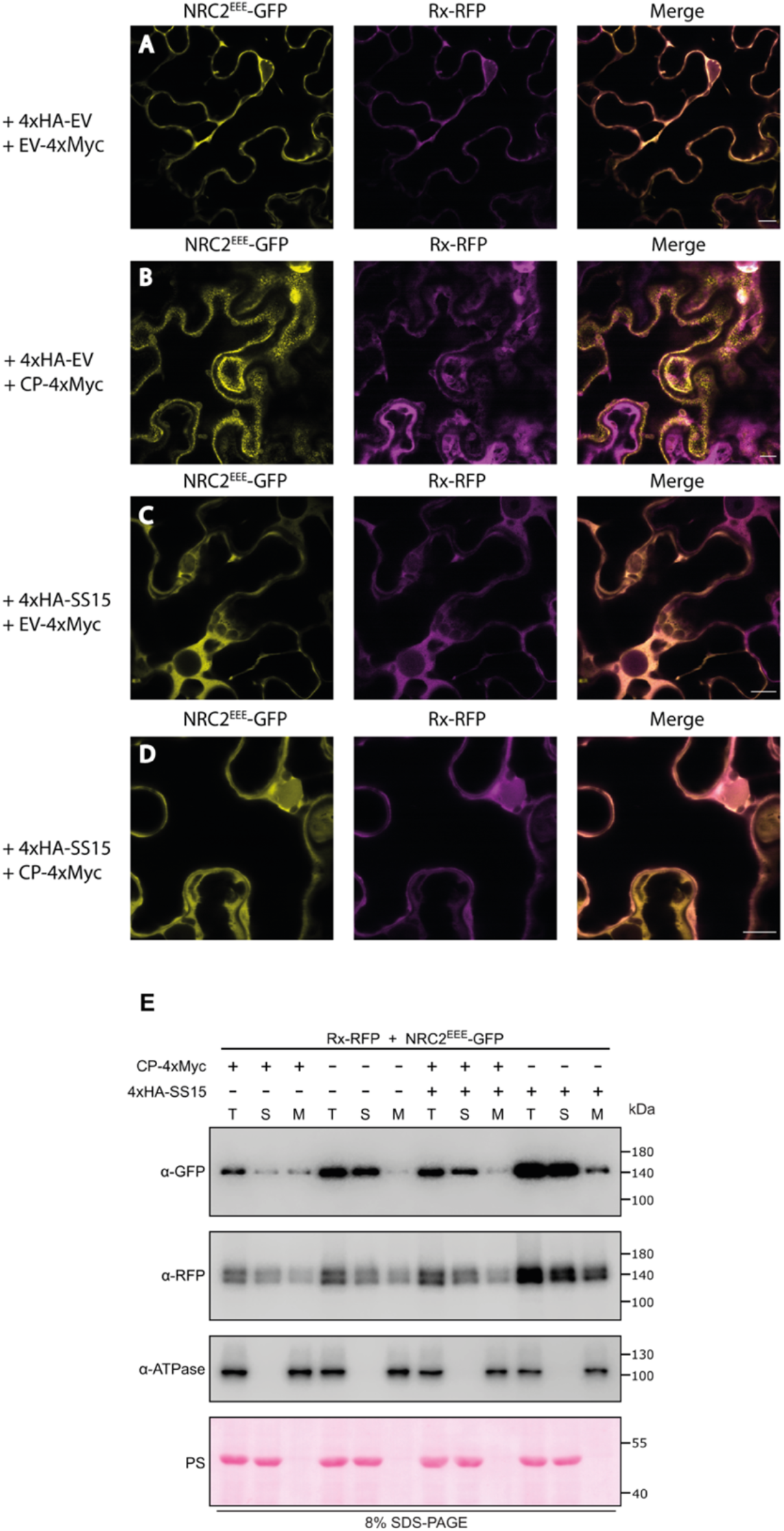
SS15 inhibits plasma membrane-association of activated NRC2. (**A-D**) C-terminally GFP-tagged NRC2^EEE^ and C-terminally RFP-tagged Rx were co-expressed with an EV-4xMyc construct or a CP-4xMyc construct in leaves of *nrc2/3/4* KO *N. benthamiana*. Representative single-plane confocal micrographs show the localization of both components of the inactive and active Rx-NRC2 system. Scale bars represent 10 µm. (**A**) NRC2^EEE^-GFP and Rx-RFP co-localize in the cytoplasm. (**B**) As reported previously, Rx/CP activated NRC2^EEE^ forms plasma membrane-associated puncta while Rx remains in the cytoplasm. (**C**) Co-expression with SS15 does not alter the localization of inactive NRC2^EEE^-GFP or Rx-RFP. (**D**) Upon co-expression CP-4xMyc, Rx-RFP and NRC2^EEE^-GFP with SS15, the punctate localization for NRC2^EEE^-GFP is no longer observed. (**E**) Membrane enrichment assays are consistent with microscopy. As reported previously, inactive NRC2^EEE^-GFP is mostly present in the soluble fraction (S) whereas activated NRC2^EEE^-GFP exhibits equal distribution across soluble and membrane (M) fractions. Upon co-expression with SS15, NRC2^EEE^-GFP distribution remains in the soluble fraction regardless of the presence or absence of PVX CP. The experiment was repeated twice with similar results.

**Fig. S3:**
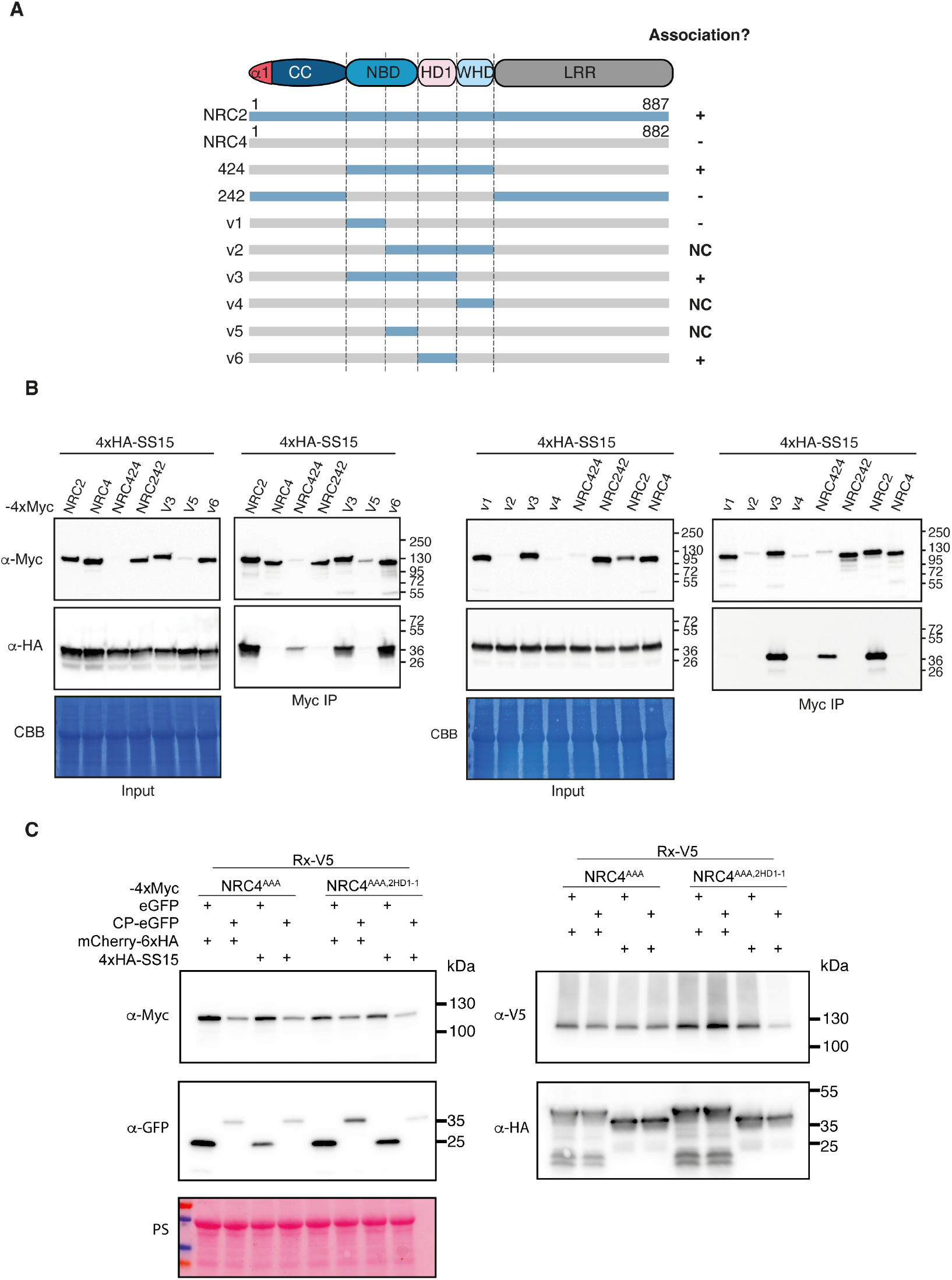
SS15 inhibits NRC2 by interacting with the HD1-1 region of the NB-ARC domain. **(A)** Schematic representation of all NRC2-NRC4 chimeric proteins generated. Association with SS15 (+) or lack thereof (-) is indicated on the right. (**B**) Co-Immunoprecipitation (Co-IP) assays between SS15 and chimeric NRC2-NRC4 variants. C-terminally 4xMyc-tagged NRC proteins were transiently co-expressed with N-terminally 4xHA-tagged SS15. IPs were performed with agarose beads conjugated to Myc antibodies (Myc IP). Total protein extracts were immunoblotted with the appropriate antisera labelled on the left. Approximate molecular weights (kDa) of the proteins are shown on the right. Rubisco loading control was carried out using Ponceau stain (PS). The experiment was repeated three times with similar results. (**C**) SDS-PAGE accompanying BN-PAGE shown in **Fig. 2E**. Total protein extracts were immunoblotted with the appropriate antisera labelled on the left. Approximate molecular weights (kDa) of the proteins are shown on the right. Rubisco loading control was carried out using Ponceau stain (PS). The experiment was repeated three times with similar results.

**Fig. S4:**
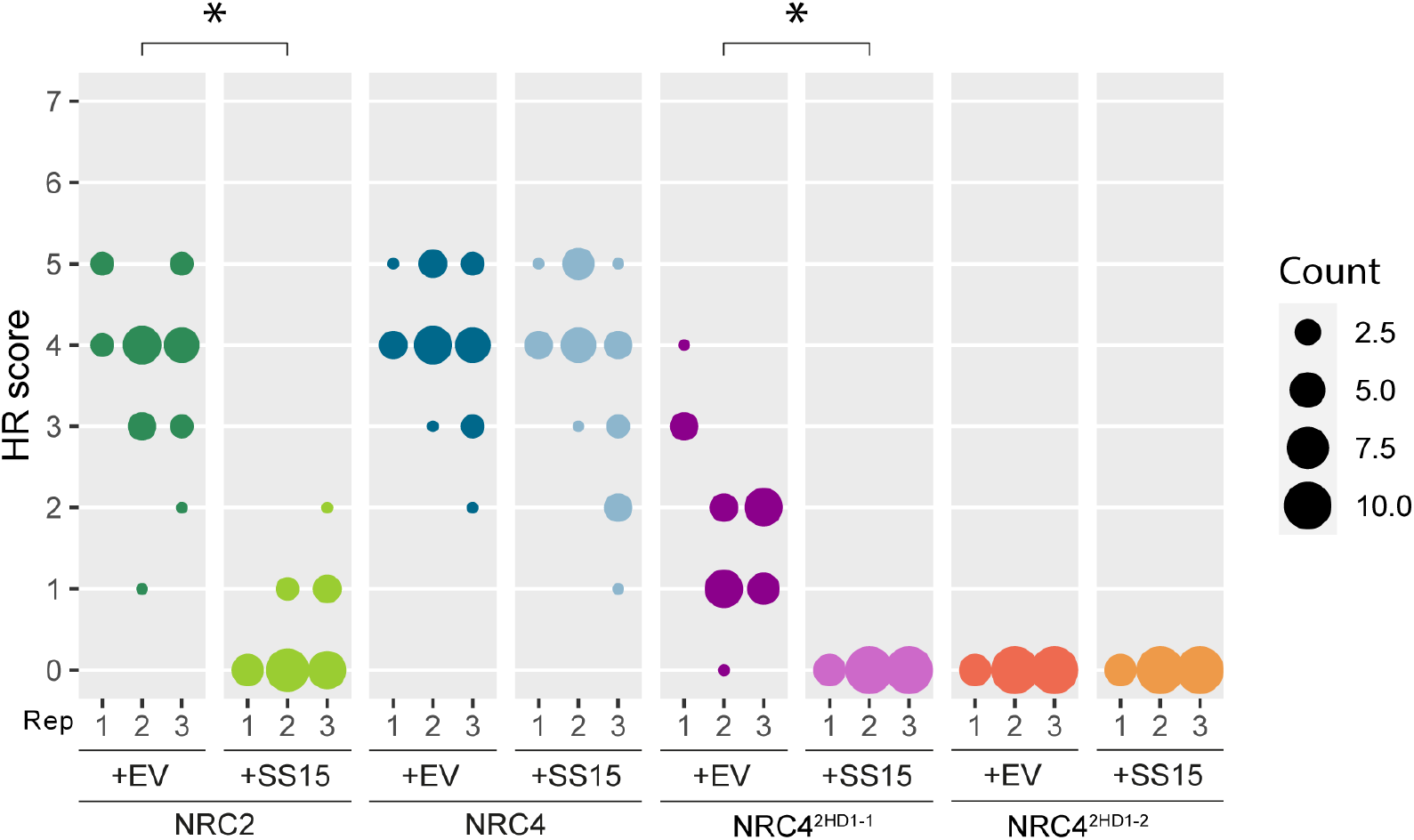
NRC4^HD1-1^ chimera is susceptible to inhibition by SS15. HR scores accompanying **Fig. 2D**. In all cases, Rx/CP was used to activate the system. HR was scored based on a modified 0–7 scale (*49*) between 5–7 days post-infiltration. HR scores are presented as dot plots, where the size of each dot is proportional to the number of samples with the same score (Count). Results are based on 3 biological replicates. Statistical tests were implemented using the besthr R library (*50*). We performed bootstrap resampling tests using a lower significance cut-off of 0.025 and an upper cut-off of 0.975. Mean ranks of test samples falling outside of these cut-offs in the control samples bootstrap population were considered significant. Significant differences between the conditions are indicated with an asterisk (*).

**Fig. S5:**
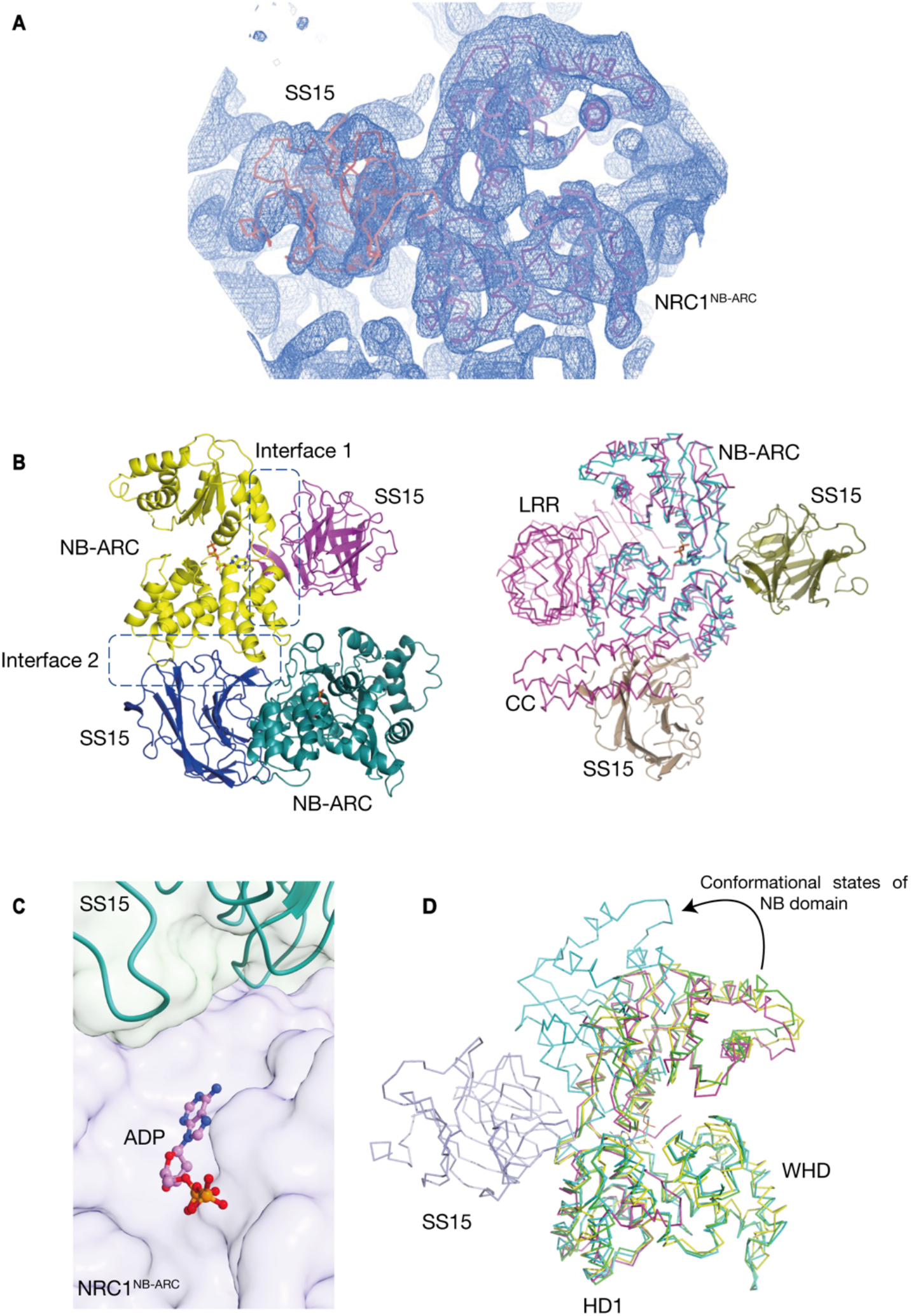
Crystal structure of SS15 in complex with SlNRC1^NB-ARC^. Electron density map showing the relative orientation and arrangement of SS15 (orange) and SlNRC1^NB-ARC^ (violet) within an asymmetric unit. 2Fo-Fc map countered at 1σ **(B)** Two possible interfaces between SS15 and SlNRC1^NB-ARC^ revealed from the crystal packing. Both interfaces (Interface 1 and Interface 2) are outlined (Left). Modelling of both potential binding interfaces for SS15 complex with full length SlNRC1 (magenta) reveals a steric clash between the CC-domain of SlNRC1 and SS15, making interface 2 unlikely to be biologically relevant in the full-length context (Right). **(C)** Close up view of interaction between SS15-SlNRC1^NB-ARC^ interaction interface relative to the ATP-binding site within the NB-ARC domain of SlNRC1. The pyrophosphate moiety of ADP is oriented facing opposite the SS15 binding interface (shown as ball and sticks), suggesting that SS15 is unlikely to displace bound nucleotide or prevent ATP hydrolysis. **(D)** Structure of SS15-SlNRC1^NB-ARC^ (yellow, PDB 8BV0) is superimposed over the NB-ARC domain of AtZAR1 in its inactive (green, PDB 6J5W), intermediate (cyan, PDB 6J5V), and active resistosome (magenta, 6J5T) conformations. Visualizing these three states reveals the trajectory of the NB domain as it moves relative to the HD1 and WHD domains while changing from inactive to activated states. The binding of SS15 at the critical hinge region between the NB and HD1-WHD domains likely immobilizes this loop, preventing these critical intramolecular rearrangements and therefore preventing NLR activation. See **Movie S1**.

**Fig. S6:**
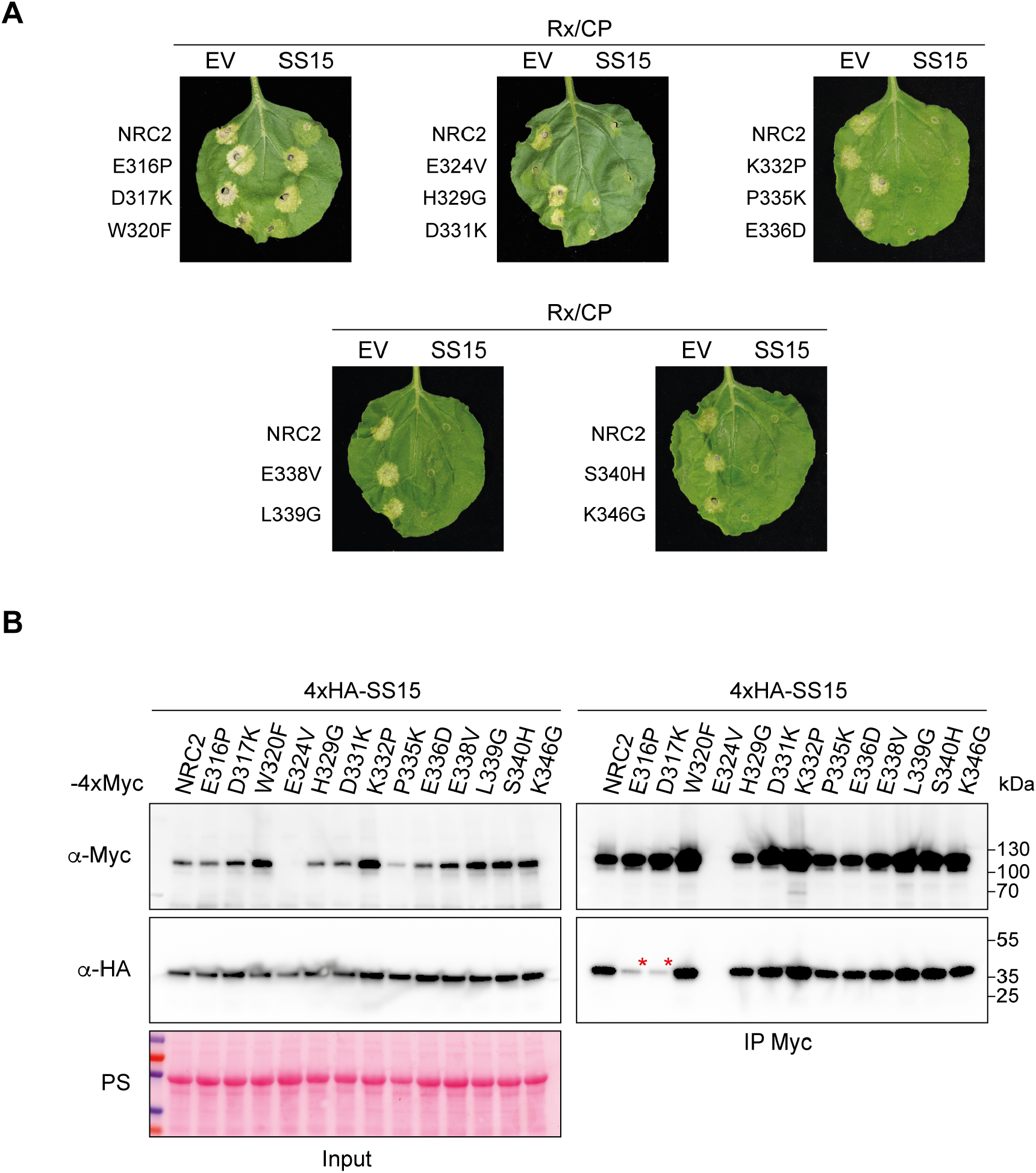
Out of 13 NRC2 variants tested, only E316P and D317K mutations abolish SS15 association and HR suppression. **(A)** Photo of representative leaves from *N. benthamiana nrc2/3/4* KO plants showing HR after co-expression of Rx and PVX CP with NRC2, or the different NRC2 variants generated. These effector-sensor-helper combinations were co-expressed with a free mCherry-6xHA fusion protein (EV) or with N-terminally 4xHA-tagged SS15. (**B**) Co-Immunoprecipitation (Co-IP) assays between SS15 and NRC2 variants. C-terminally 4xMyc-tagged NRC2 variants were transiently co-expressed with N-terminally 4xHA-tagged SS15. IPs were performed with agarose beads conjugated to Myc antibodies (Myc IP). Total protein extracts were immunoblotted with appropriate antisera labelled on the left. Approximate molecular weights (kDa) of the proteins are shown on the right. Rubisco loading control was carried out using Ponceau stain (PS). The experiment was repeated three times with similar results.

**Fig. S7:**
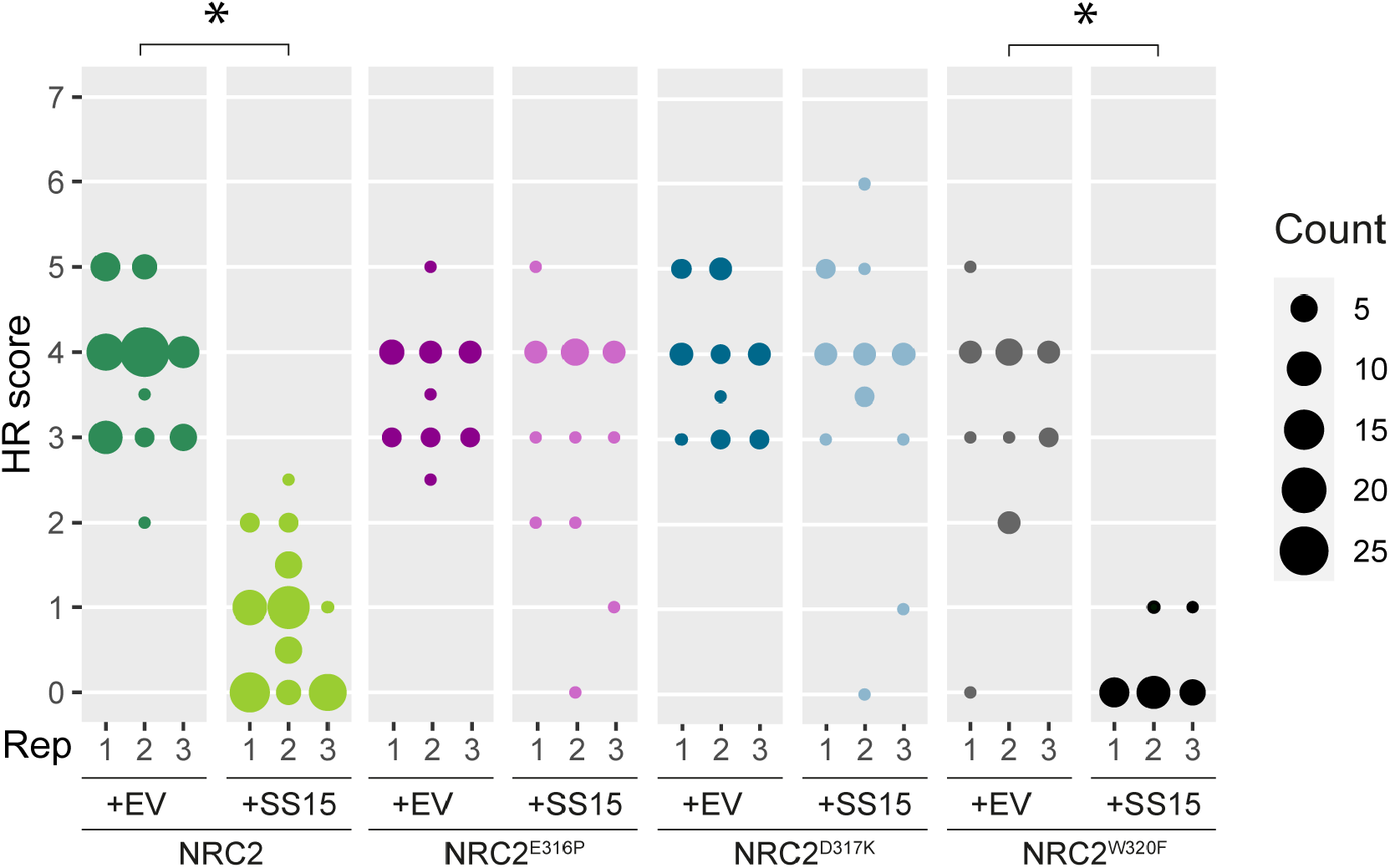
NRC2^E316P^ and NRC2^D317K^ abolish SS15-mediated suppression of Rx. HR scores accompanying **Fig. 3C**. In all cases, Rx/CP was used to activate the system. HR was scored based on a modified 0–7 scale (*49*) between 5–7 days post-infiltration. HR scores are presented as dot plots, where the size of each dot is proportional to the number of samples with the same score (Count). Results are based on 3 biological replicates. Statistical tests were implemented using the besthr R library (*50*). We performed bootstrap resampling tests using a lower significance cut-off of 0.025 and an upper cut-off of 0.975. Mean ranks of test samples falling outside of these cut-offs in the control samples bootstrap population were considered significant. Significant differences between the conditions are indicated with an asterisk (*).

**Fig. S8:**
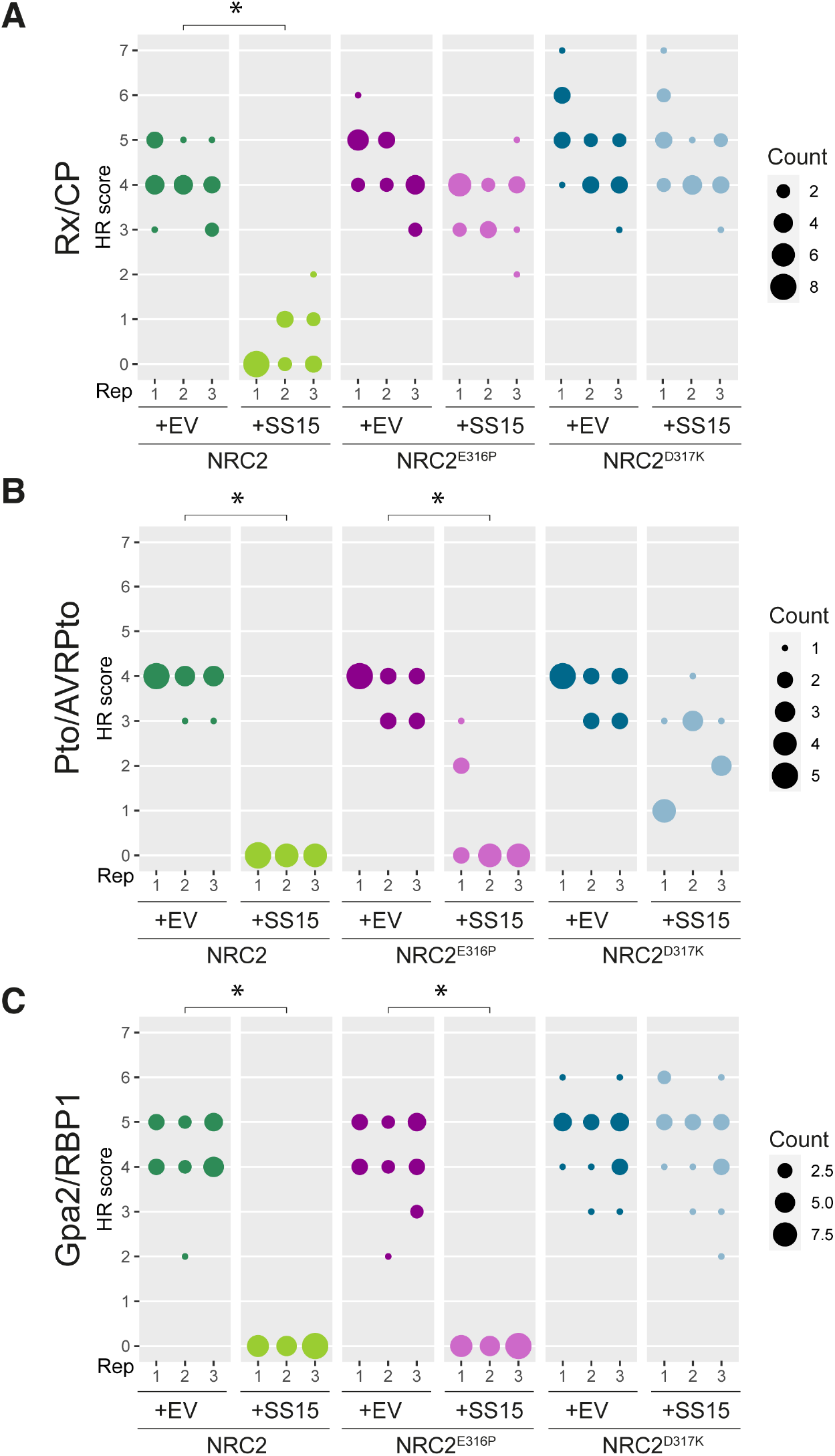
NRC2^D317K^ abolishes SS15-mediated suppression of Rx, Gpa2 and Prf. HR scores accompanying **Fig. 4A**. NRCs were activated using Rx/CP (**A**), Pto/AVRPto (**B**) or Gpa2/RBP1 (**C**). HR was scored based on a modified 0–7 scale (*49*) between 5–7 days post-infiltration. HR scores are presented as dot plots, where the size of each dot is proportional to the number of samples with the same score (count). Results are based on 3 biological replicates. Statistical tests were implemented using the besthr R library (*50*). We performed bootstrap resampling tests using a lower significance cut-off of 0.025 and an upper cut-off of 0.975. Mean ranks of test samples falling outside of these cut-offs in the control samples bootstrap population were considered significant. <u>Significant differences between the conditions are indicated with an asterisk (*)</u>.

**Fig. S9:**
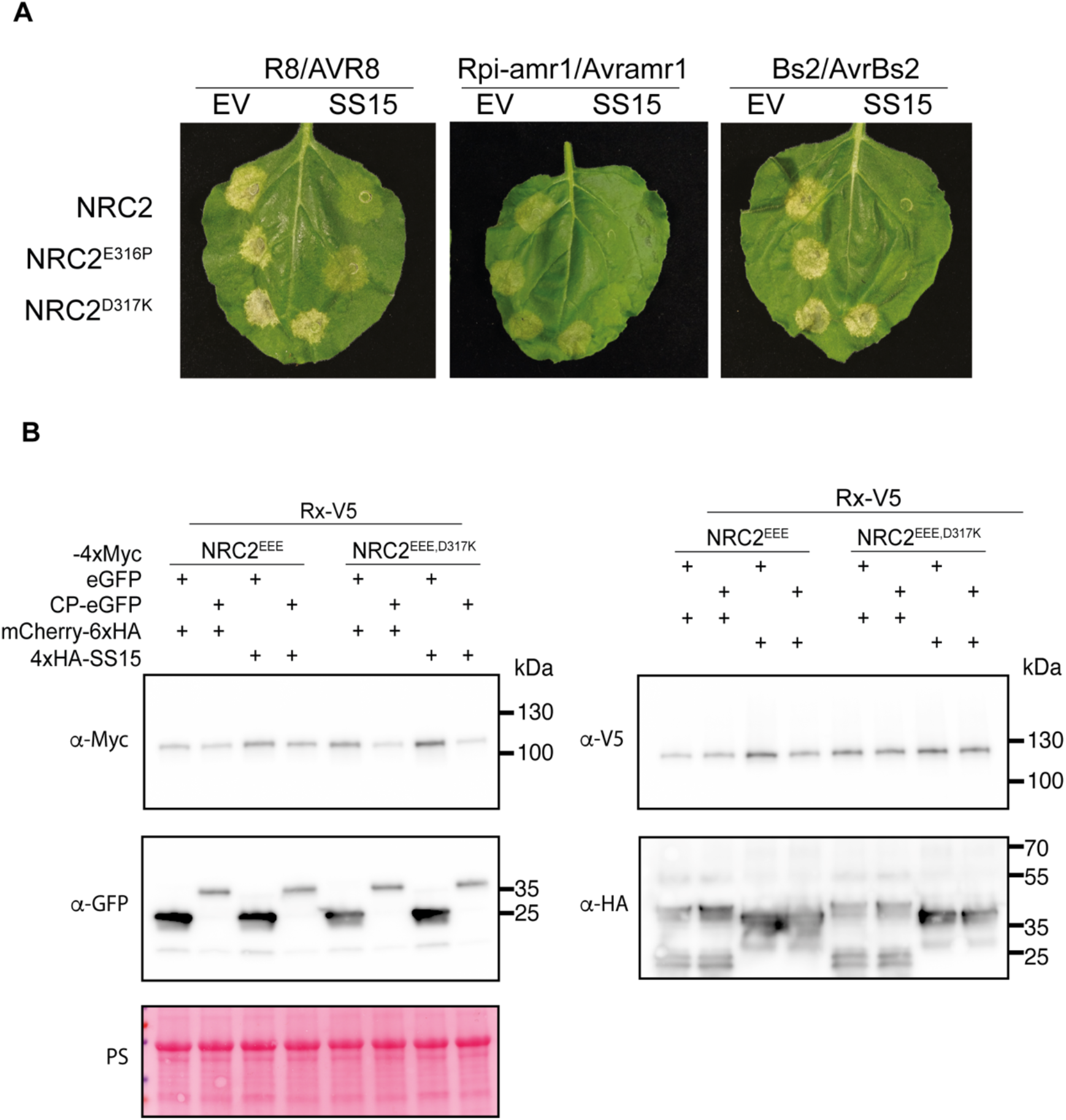
NRC2^D317K^ abolishes SS15-mediated suppression of all NRC2-dependent sensors tested and restores NRC2 resistosome formation. Photo of representative leaves from *N. benthamiana nrc2/3/4* KO plants showing HR after co-expression of NRC2, or different NRC2 variants generated with various sensor/effector pairs. These effector-sensor-helper combinations were co-expressed with a free mCherry-6xHA fusion protein (EV) or with N-terminally 4xHA-tagged SS15. (**B**) SDS-PAGE accompanying BN-PAGE shown in **Fig. 4B**. Total protein extracts were immunoblotted with the appropriate antisera labelled on the left. Approximate molecular weights (kDa) of the proteins are shown on the right. Rubisco loading control was carried out using Ponceau stain (PS). The experiment was repeated three times with similar results.

## Supplementary Tables

**Table S1: List of constructs used in this study**. (As separate file)

**Table S2: List of OD**_**600**_ **used for agroinfiltration experiments**. (As separate file)

**Table S3:**
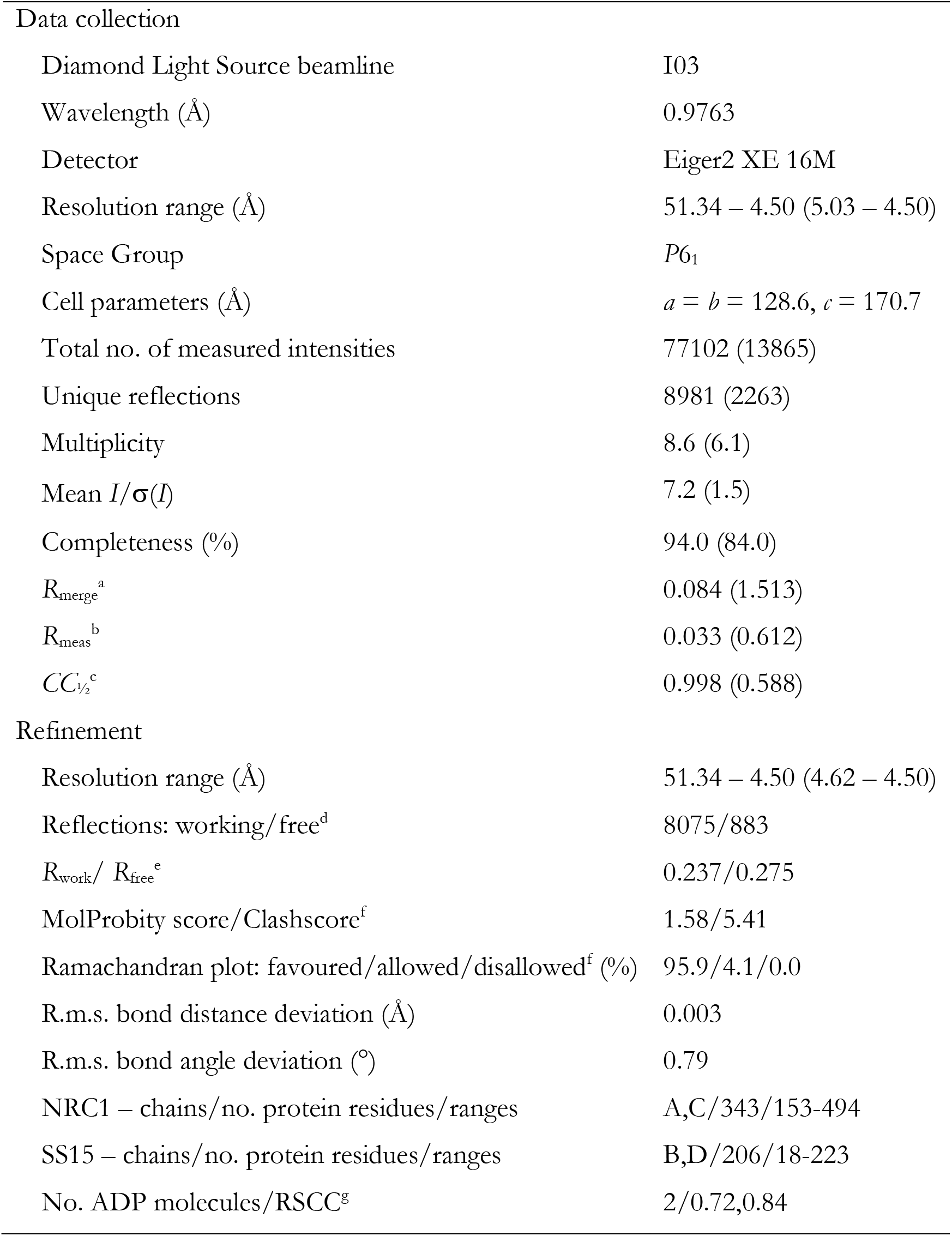
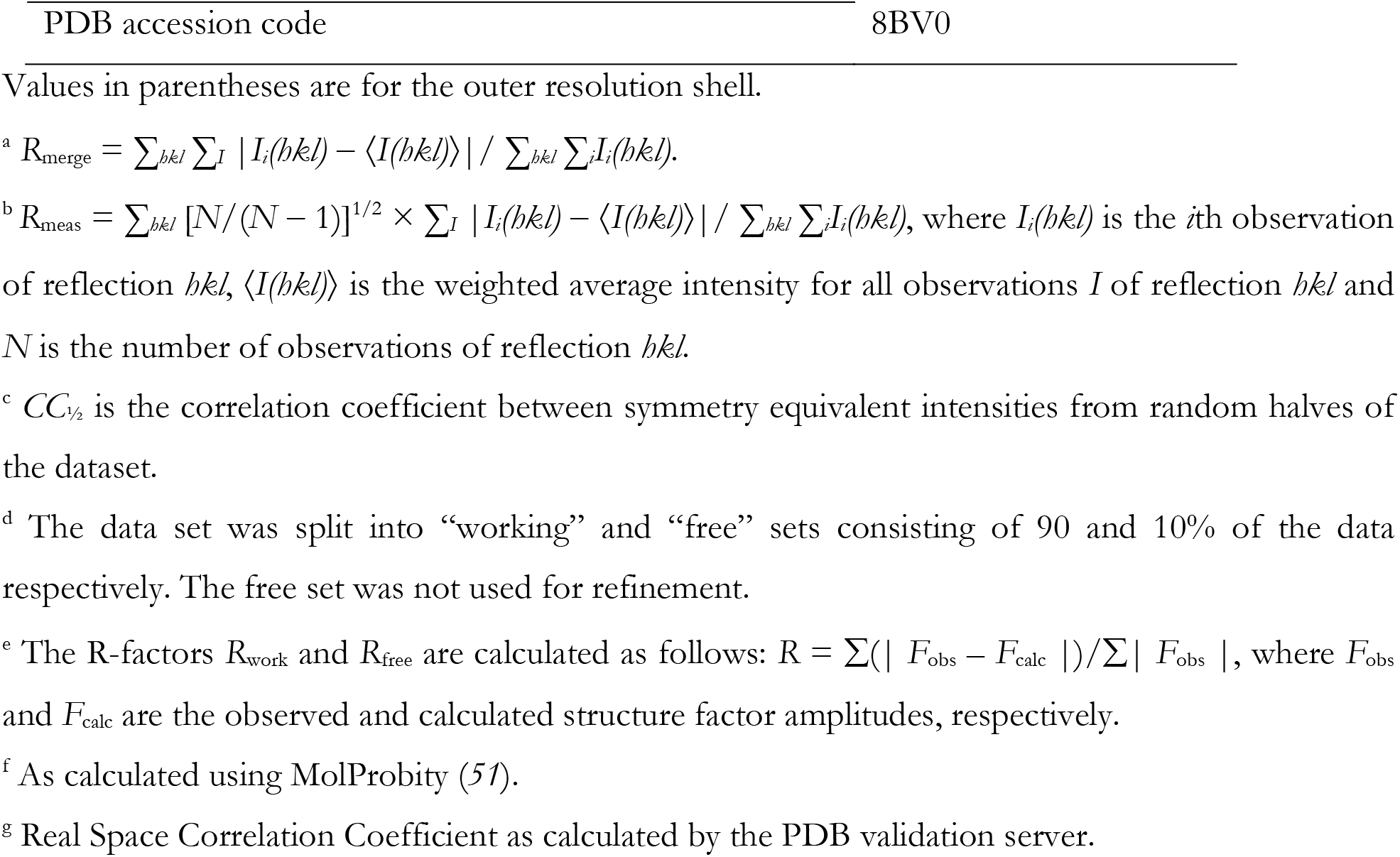
Summary of X-ray data and model parameters for NRC1-SS15.

|  |  |
| --- | --- |
| Data collection |  |
| Diamond Light Source beamline | I03 |
| Wavelength (Å) | 0.9763 |
| Detector | Eiger2 XE 16M |
| Resolution range (Å) | 51.34 – 4.50 (5.03 – 4.50) |
| Space Group | *P*6<sub>1</sub> |
| Cell parameters (Å) | *a* = *b* = 128.6, *c* = 170.7 |
| Total no. of measured intensities | 77102 (13865) |
| Unique reflections | 8981 (2263) |
| Multiplicity | 8.6 (6.1) |
| Mean *I*/σ(*I*) | 7.2 (1.5) |
| Completeness (%) | 94.0 (84.0) |
| *R*<sub>merge</sub><sup>a</sup> | 0.084 (1.513) |
| *R*<sub>meas</sub><sup>b</sup> | 0.033 (0.612) |
| *CC*<sub>1/2</sub><sup>c</sup> | 0.998 (0.588) |
| Refinement |  |
| Resolution range (Å) | 51.34 – 4.50 (4.62 – 4.50) |
| Reflections: working/free<sup>d</sup> | 8075/883 |
| *R*<sub>work</sub>/ *R*<sub>free</sub><sup>e</sup> | 0.237/0.275 |
| MolProbity score/Clashscore<sup>f</sup> | 1.58/5.41 |
| Ramachandran plot: favoured/allowed/disallowed<sup>f</sup> (%) | 95.9/4.1/0.0 |
| R.m.s. bond distance deviation (Å) | 0.003 |
| R.m.s. bond angle deviation (°) | 0.79 |
| NRC1 – chains/no. protein residues/ranges | A,C/343/153-494 |
| SS15 – chains/no. protein residues/ranges | B,D/206/18-223 |
| No. ADP molecules/RSCC<sup>g</sup> | 2/0.72,0.84 |

Values in parentheses are for the outer resolution shell.
$^b R_{\text{meas}} = \sum_{hkl} [N/(N-1)]^{1/2} \times \sum_I |I_i(hkl) - \langle I(hkl) \rangle| / \sum_{hkl} \sum_I I_i(hkl)$ , where $I_i(hkl)$ is the $i$ th observation of reflection $hkl$ , $\langle I(hkl) \rangle$ is the weighted average intensity for all observations $I$ of reflection $hkl$ and $N$ is the number of observations of reflection $hkl$ .
$^c CC_{1/2}$ is the correlation coefficient between symmetry equivalent intensities from random halves of the dataset.
$^d$ The data set was split into “working” and “free” sets consisting of 90 and 10% of the data respectively. The free set was not used for refinement.
$^e$ The R-factors $R_{\text{work}}$ and $R_{\text{free}}$ are calculated as follows: $R = \sum(|F_{\text{obs}} - F_{\text{calc}}|) / \sum |F_{\text{obs}}|$ , where $F_{\text{obs}}$ and $F_{\text{calc}}$ are the observed and calculated structure factor amplitudes, respectively.
$^f$ As calculated using MolProbity (51).
$^g$ Real Space Correlation Coefficient as calculated by the PDB validation server.

## Supplementary Movies

**Movie S1: SS15 immobilizes hinge region between NB and HD1-WHD domains of NRCs to prevent NLR activation**.

## Acknowledgements

We are very thankful to Jiorgos Kourelis, Clemence Marchal, Daniel Lüdke and Hee-Kyung Ahn (The Sainsbury Laboratory, Norwich, UK) for invaluable scientific discussions, ideas, and feedback. We thank Julia Mundy for assistance with crystallography experiments. We thank the Diamond Light Source, UK (beamline i03 under proposal MX18565) for access to X-ray data collection facilities. M.P.C. thanks Emiliano “Dibu” Martínez and Lionel Messi for inspiration. We thank all members of the TSL Support Services for their invaluable assistance.

## Author Contributions

Conceptualization: M.P.C, C-H.W., S.K., L.D.

Methodology: M.P.C., H.P., M.S., D.M.L., C.E.M.S., L.D.

Validation: M.P.C., H.P., M.S., Y.T., C.D., E.L.H.Y., A.H.

Formal Analysis: M.P.C., H.P., M.S., A.T., D.M.L., Y.T., C.D., C.E.M.S.

Investigation: M.P.C., H.P., M.S., Y.T., C.D., E.L.H.Y., A.H., L.D.

Data Curation: M.P.C., H.P., M.S., A.T., D.M.L., Y.T., C.D., E.L.H.Y., C.E.M.S., A.H.

Visualization: M.P.C., H.P., M.S., A.T., Y.T., C.D., E.L.H.Y., L.D.

Writing – Original Draft: M.P.C., S.K., L.D.

Writing – Review & Editing: M.P.C., D.M.L., S.K., L.D.

Supervision: M.P.C., T.O.B., S.K., L.D.

Project Administration: M.P.C., T.O.B., S.K., L.D.

Funding Acquisition: T.O.B., S.K., L.D.

## Data and materials availability

All relevant data are within the paper, in the Supplementary Materials and Source Data files. Data relevant to the structure presented in **Fig. 3** can be found in the Protein Data Bank, PDB 8BV0.

